# Stimulus dependent relationships between behavioral choice and sensory neural responses

**DOI:** 10.1101/2019.12.27.889550

**Authors:** Daniel Chicharro, Stefano Panzeri, Ralf M. Haefner

## Abstract

Understanding the relationship between trial-to-trial variability in neural responses of sensory areas and behavioral choices is fundamental to elucidate the mechanisms of perceptual decision-making. In two-choice tasks, activity-choice co-variations have traditionally been quantified with choice probabilities (CP). It has been so far commonly assumed that choice-related neural signals are separable from stimulus-driven responses, which has led to characterizing activity-choice covariations only with a single CP value estimated combining trials from all stimulus levels. In this work we provide theoretical and experimental evidence for the stimulus dependence of the relationship between neural responses and behavioral choices. We derived a general analytical CP expression for this dependency under the general assumption that a decision threshold converts an internal stimulus estimate into a binary choice. This expression predicts a stereotyped threshold-induced CP modulation by the stimulus information content. We reanalyzed data from Britten et al. (1996) and found evidence of this modulation in the responses of macaque MT cells during a random dot discrimination task. Moreover, we developed new methods of analysis that allowed us to further identify a richer structure of cell-specific CP stimulus dependencies. Finally, we capitalised on this progress to develop new generalized linear models (GLMs) with stimulus-choice interaction terms, which show a higher predictive power and lead to a more precise assessment of how much each neuron is stimulus- or choice-driven, hence allowing a more accurate comparison across areas or cell types. Our work suggests that characterizing the patterns of stimulus dependence of choice-related signals is essential to properly determine how neurons in different areas contribute to linking sensory representations to perceptual decisions.

## 1 Introduction

How perceptual decisions depend on responses of sensory neurons is a fundamental question in systems neuroscience (Parker and Newsome, 1998; Gold and Shadlen, 2001; Romo and Salinas, 2003; Gold and Shadlen, 2007; Siegel et al., 2015; van Vugt et al., 2018; O’Connell et al., 2018; Steinmetz et al., 2019). The seminal work of Britten et al. (1996) showed that responses from single cells in area MT of monkeys during the performance of a random dot discrimination task covaried with behavioral choices. Similar activity-choice covariations have been found in multiple sensory areas and for a variety of two-choice tasks, including both discrimination and detection tasks (see Nienborg et al., 2012; Cumming and Nienborg, 2016, for a review). Identifying the location of cells whose activity encodes choice and how and when choice information is encoded in neural activity is essential to understand how the brain generates appropriate behaviors based on sensory information.

In the context of two-choice tasks, Choice Probabilities (CP) have been the most prominent measure (Britten et al., 1996; Parker and Newsome, 1998; Nienborg et al., 2012) used to quantify activity-choice covariations. Most commonly, it is assumed that choice-related neural signals are independent of and separable from stimulus-driven responses, and a single CP value is calculated per cell, which quantifies the global strength of activity-choice covariations (so-called grand CP (Britten et al., 1996)). This assumption was originally motivated by the work of Britten et al. (1996), who found no significant dependence of CPs on the coherence level of the random dots presented in the discrimination task. Similarly, when activity-choice covariations are not quantified alone as in the CP, but modeled jointly with other covariates of the neural responses using Generalized Linear Models (GLMs) (Truccolo et al., 2005), the stimulus level and the choice value are also usually used as separate predictors of the responses (Park et al., 2014; Runyan et al., 2017).

Theoretically, the interpretation of choice-related signals has been guided by computational and analytical results derived from a model of decision making in which a continuous internal estimate of the stimulus is converted by a threshold mechanism into a behavioral choice (Shadlen et al., 1996; Cohen and Newsome, 2009). Based on this feedforward model, Haefner et al. (2013) derived an analytical expression of the CP measure which explains the role originating the activity-choice covariations of the read-out weights and of correlated trial-to-trial variability across the population of sensory neurons. However, this expression was derived only for non-informative stimuli and hence does not address how the stimulus content can modulate choice-related signals when the choice is estimated from sensory responses through a threshold mechanism. Furthermore, while this model provides an interpretation of the CP based on feedforward mechanisms, it remains an unresolved question to determine to which degree feedforward or feed-back signals account for activity-choice covariations in different areas and stages of the perceptual decision making process (Cumming and Nienborg, 2016). In particular, the structure of decision-related feedback signals (Bondy et al., 2018) is expected to induce cell-specific modulations of choice-related signals by the stimulus content.

The interaction between stimulus and choice signals in neural activity has largely been unexplored. However, if the assumption that these signals are independent and separable is incorrect, then previously reported quantifications of whether, and how much, each area is stimulus- or choice-driven may need to be re-evaluated. In this work, we reassess the existence of stimulus dependencies in activity-choice covariations by making progress both with models and with neural data analysis tools.

First, we extended the analytical model of CP of Haefner et al. (2013) to the general case of informative stimuli and including both a feedforward or feedback origin of the covariation between the choice and each cell. This generalized model predicts a stimulus dependence of CP induced by the decision threshold, consisting in a symmetric increase of the magnitude of the CP for more informative stimuli. This increase is multiplicative and of opposite sign for cells with opposite choice preferences, which indicates that an analysis of CP-stimulus dependencies at the population level (Britten et al., 1996) may not detect it. Second, we developed new analytical methods that increase the power to detect CP-stimulus dependencies analyzing within-cell patterns of dependence. Traditionally, grand CPs are estimated pooling trials from all stimulus levels, after discounting the estimated stimulus-driven component from the responses to informative stimuli (Nienborg and Cumming, 2009; Kang and Maunsell, 2012). On the other hand, our new methods offer a solution to the methodological challenge posed by the fact that highly informative stimuli lead to fewer samples of responses for the error choice, and allow characterizing a profile of CP values as a function of stimulus levels.

We applied to the MT cells dataset of Britten et al. (1996) our combination of model-driven analyses and finer methods to unravel stimulus dependencies of CPs previously overlooked by traditional analyses of the same dataset. The patterns of CP-stimulus dependencies that we found provide evidence for the symmetric threshold-induced dependence predicted by our CP model, but also reveal a richer structure of dependencies. The main features of this structure consist in the existence of asymmetric patterns with higher CPs for stimuli eliciting higher activity and a coupling between the average CP strength and the degree of pattern asymmetry. We show that these features are consistent with the effect of excitability fluctuations (Goris et al., 2014) in the responses of the sensory neurons contributing to the perceptual decision.

Moreover, we also demonstrate the utility of introducing new stimulus-choice interaction terms in Generalized Linear Models (GLMs). GLMs containing interaction terms improve the fitting of the responses of the MT cells and, importantly, affect the quantification of how stimulus-driven versus choice-driven each cell is. Because the importance of these terms depends itself on cell-specific properties, incorporating them is expected to refine population-level comparisons, e.g. across cell types, areas, or tuning properties. Together, our advances lay down the basis for a better understanding of the distribution of stimulus and choice signals within each neuron and across brain areas and neural populations.

## 2 Results

We will first present the theoretical analysis of choice probabilities, followed by the analysis of the data from Britten et al. (1996) applying our new methods to quantify stimulus-dependent activity-choice covariations with CPs and GLMs.

### 2.1 Choice probabilities with informative stimuli

Decision-related signals, independent of their origin, lead to activity-choice co-variations between the neural response of a cell *i, r*_*i*_, and the psychophysical choice *D*. For tasks involving two choices (both discrimination and detection tasks), this covariation is captured by the difference between the neural response distributions conditioned on the behavioral choice, *p*(*r*|*D* = −1) and *p*(*r*|*D* = 1). The choice probability (Britten et al., 1996) quantifies the difference between these two distributions:

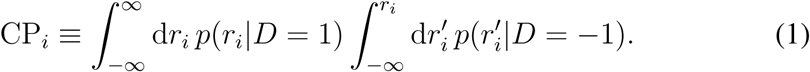

Intuitively, the CP is defined as the probability that a random sample from the distribution for choice 1 trials is larger than a random sample from the distribution for choice −1 trials (Britten et al., 1996; Parker and Newsome, 1998; Nienborg et al., 2012). It is 0.5 if both distributions are identical and goes to 0 or 1 as they are more and more separated. While the CP quantifies the activity-choice covariations without any assumption about the decision-making mechanisms (Fig. 1A), the interpretation of CP values has been informed by the conceptualization of threshold models (Shadlen et al., 1996) in which a decision variable *d*, corresponding to an internal estimate of the stimulus, determines the choice by comparing *d* to a threshold *θ*. The internal estimate is usually modeled as a weighted sum of the neural responses. Based on this model, Haefner et al. (2013) derived a CP expression for the case of a purely feedforward threshold model and uninformative stimuli.

**Figure 1:**
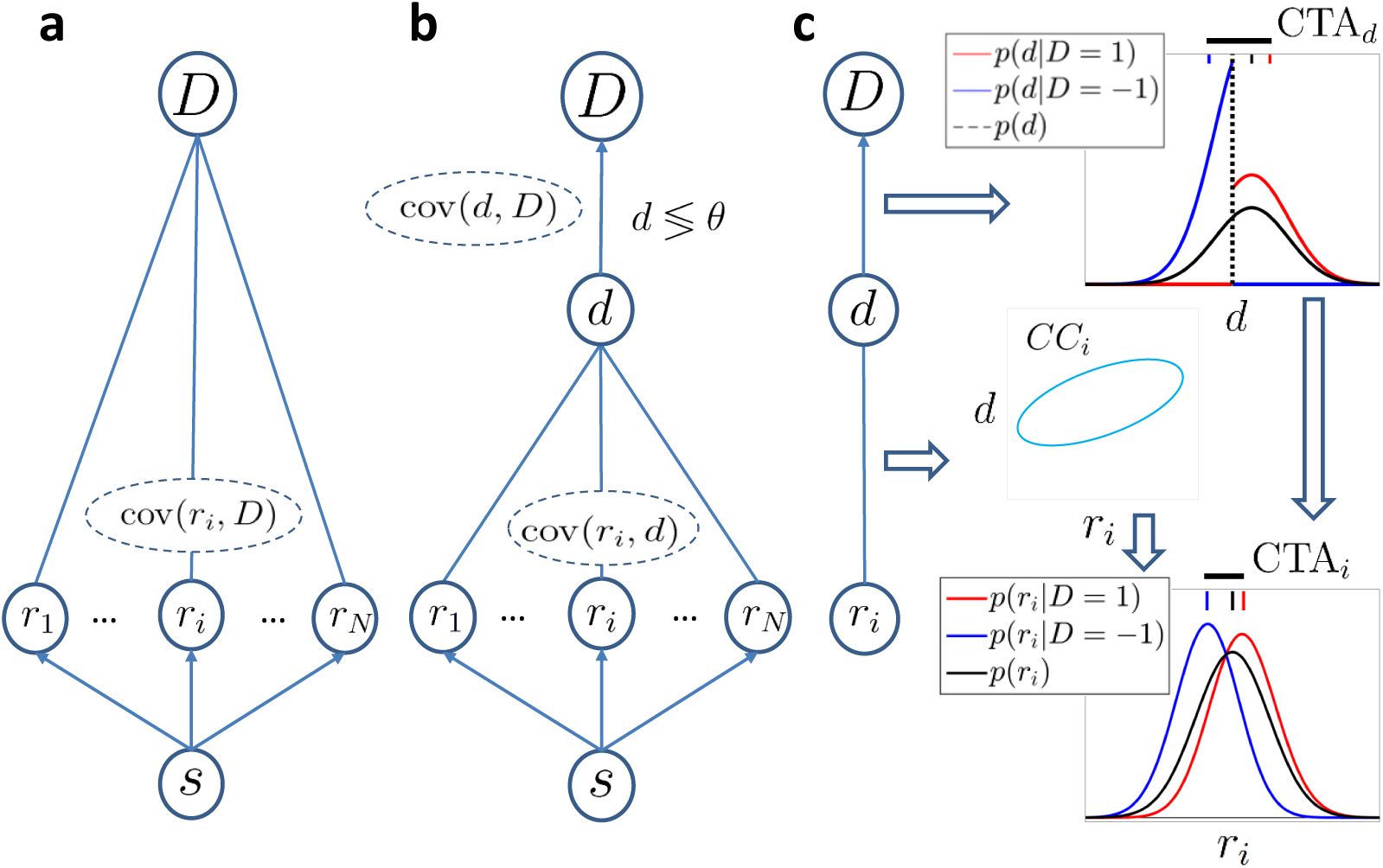
Models of choice probabilities. Arrows indicate causal influences. Undirected edges indicate a relationship that may be due to feedforward and/or feedback signals. **a)** A model agnostic to the causal origin of the choice–response covariation: Sensory neurons encode a stimulus *s* and activity covaries with choice *D*. **b)** Threshold model with a continuous decision variable *d* that mediates between responses and the choice. The decision is made comparing *d* with a threshold *θ*. **c)** Signature of the threshold mechanism into activity-choice covariations. The threshold mechanism (vertical dashed black line) dichotomizes the space of *d*, resulting in a difference between the mean of the conditional distributions associated with *D* = ±1 (red and blue vertical top bars, respectively). This difference is quantified by CTA_*d*_ (horizontal thick black line) and propagates to the Choice-Triggered Average (CTA_*i*_) of the responses when some correlation CC_*i*_ exists between *d* and *r*_*i*_.

We here extended the derivation in Haefner et al. (2013) to obtain a general expression of the CP to analyze the effect of informative stimuli regardless of the feedforward or feedback origin of the dependencies between the neural responses and the decision variable (Fig. 1B). The exact solution of this model is described in the Methods section (Eq. 14). Here we focus on a linear approximation derived from the new general solution, which is accurate for a very wide range of informative stimuli (see Methods) and captures the threshold-induced dependence of the CP on the stimulus content. In particular, the linear approximation expresses the CP_*i*_ of neuron *i* as

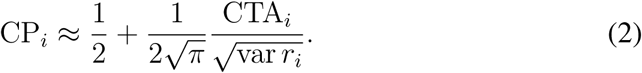

where var *r*_*i*_ is the variance of the responses of cell *i* and CTA_*i*_ is its Choice-Triggered Average, defined as the difference between the mean response for trials of each choice (Haefner, 2015; Chicharro et al., 2017)

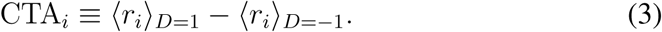

The CTA generically quantifies the linear dependencies between the responses and choice, and hence the approximation of the CP does not depend on their feed-forward or feedback origin (Fig. 1A).

In the threshold model, the mediating continuous decision variable *d* allows breaking down the covariance cov(*r*_*i*_, *D*) in terms of the relationships between *r* and *d*, and between *d* and *D*, respectively (Fig. 1B). The correlation coefficient between activity and choice is factorized such that corr(*r*_*i*_, *D*) = corr(*r*_*i*_, *d*)corr(*d, D*).

Furthermore, given the binary nature of choice *D*, the CTA is directly proportional to the covariance of the responses and the choice: CTA_*i*_ = 2cov(*r*_*i*_, *D*)*p*(*D* = 1)*p*(*D* = −1) ^1^. Accordingly,

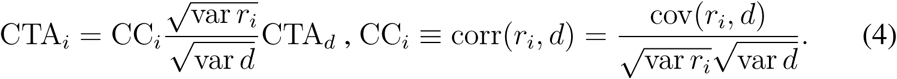

Here CC_*i*_ is the correlation coefficient between the sensory responses and the decision variable, termed choice correlation (Pitkow et al., 2015). CTA_*d*_ is the difference of the means of the (unobserved) decision variable for the two choices. Eq. 4 describes how activity-choice covariations appear in the threshold model (Fig. 1C): the threshold mechanism dichotomizes the space of the decision variable, resulting in a different mean of *d* for each choice, which is quantified in CTA_*d*_. The CTA_*d*_ is reflected for each cell into a specific CTA_*i*_, depending on the correlation CC_*i*_ between its activity *r*_*i*_ and the decision variable *d*. The dependence between the responses and *d* is quantified in CC_*i*_, while the dependence between the internal decision variable and the choice *D* is captured in 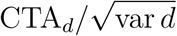.

Both of these factors of the CTA_*i*_ can be stimulus dependent. The Choice correlation CC_*i*_ may change with the stimulus due to stimulus-dependent noise correlations (Ponce-Alvarez et al., 2013) or stimulus-dependent feedback (Bondy et al., 2018), which are cell specific. Conversely, for the CTA_*d*_ the stimulus dependence is intrinsic to the effect of the threshold. This is because, analogously to the CTA_*i*_, also CTA_*d*_ = 2cov(*d, D*)*p*(*D* = 1)*p*(*D* = −1). The stimulus information alters *p*(*D* = 1)*p*(*D* = −1), and hence changes CTA_*d*_. In more detail, an informative stimulus *s* will shift the mean of *d*, thus altering the dichotomization of *d* produced by the threshold *θ* (Fig. 1C). The exact form of CTA_*d*_ depends on the distribution of the decision variable *p*(*d*). However, since the decision variable is determined by the responses of a whole population of neurons, its distribution is expected to be well approximated by a Gaussian distribution, even if the distribution of neural responses for any single neuron is not Gaussian. With this Gaussian approximation, the dichotomization of the space of *d* is specified by the probability of choosing choice 1, *p*_CR_ ≡ *p*(*D* = 1) = *p*(*d* > *θ*), which determines the ratio of the choices, i.e., the rate of each choice over trials (‘choice ratio’ or ‘choice rate’, respectively) and we can express 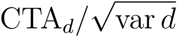 in terms of *p*_CR_. In particular, we define a factor *h*(*p*_CR_) as the ratio between 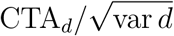 for each *p*_CR_ with respect to its value for the uninformative stimulus (*p*_CR_ = 0.5). Under gaussianity

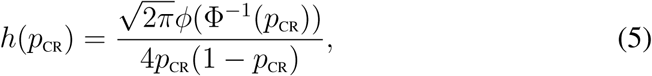

where *ϕ*(*x*) is the density function of a zero-mean, unit variance, Gaussian distribution, and Φ^−1^ is the corresponding inverse cumulative density function. By construction, *h*(*p*_CR_) = 1 for *p*_CR_ = 0.5. Finally, combining Eq. 5 with Eqs. 2 and 4, the CP is expressed as

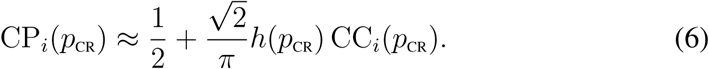

For an uninformative stimulus this CP expression corresponds to the linear approximation derived in Haefner et al. (2013). For different stimulus levels, the CP can change in two ways. First, the choice correlation CC_*i*_ itself can be stimulus dependent. Second, the multiplicative factor *h*(*p*_CR_) modulates the CP because different stimulus levels change the ratio of choices (Eq. 5). The factor *h*(*p*_CR_) only captures the covariation between *d* and *D*, and hence is common to all cells, that is, the CP_*i*_ depends on the specific response properties of cell *i* only through CC_*i*_. Furthermore, *h*(*p*_CR_) affects the CP_*i*_ independently of the source of the covariation cov(*r*_*i*_, *d*), which can be due to feedforward or feedback signals.

We now examine in more detail the shape of *h*(*p*_CR_). Fig. 2A shows CPs as a function of *p*_CR_. To characterize the shape of *h*(*p*_CR_), we set the CC to be invariant with the ratio of choices (i.e. stimulus-independent). Therefore, the CP directly reflects the dependence of *h*(*p*_CR_) on *p*_CR_. For *p*_CR_ ≠ 0.5, *h*(*p*_CR_) > 1 and the CP increases symmetrically with *p*_CR_ departing from 0.5. This can be understood for example considering feedforward contributions of neural responses to the decision variable. A highly informative stimulus induces signal-dominated neural responses, so that *d* most likely lies on the side of the threshold compatible with the sensory stimulus presented (e.g. *D* = 1) and leads to a *p*_CR_ = *p*(*D* = 1) close to 1. This means that *p*(*r*_*i*_ |*D* = 1) is similar to *p*(*r*_*i*_). On the other hand, the opposite choice is made only for trials with a substantial and contradictory departure of the neural responses from the signal-driven mean response. Accordingly, the distribution *p*(*r*_*i*_|*D* = −1) contains responses that lie in the tail of *p*(*r*_*i*_). Hence, as *p*_CR_ approaches 1, the mean of *p*(*r*_*i*_|*D* = 1) converges to the unconditional mean, while the departure of the mean of *p*(*r*_*i*_|*D* = −1) from the unconditional mean becomes increasingly large, resulting in an increasing difference between the two (Fig. 1C, and Eq. 17 in Methods).

**Figure 2:**
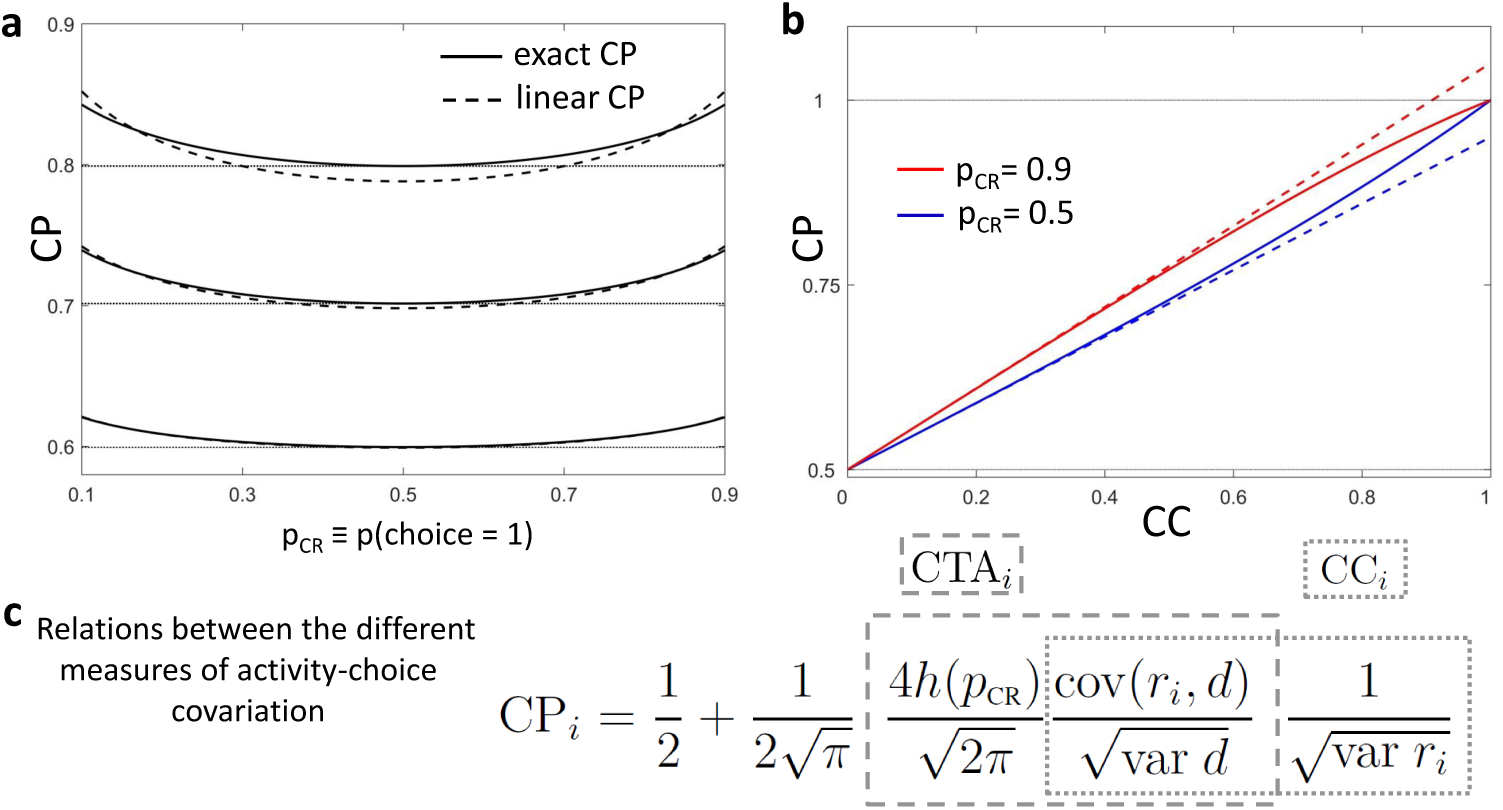
Choice probabilities calculated from the threshold model in the presence of informative stimuli. **a)** CP as a function of *p*_CR_. Results are shown for three values of a stimulus-independent choice correlation CC_*i*_. Solid lines represent the exact solution of the CP obtained from our model (see Methods, Eq. 14) and dashed lines its linear approximation (Eq. 6). **b)** Comparison of the exact solution of the CP (solid) and its linear approximation (dashed), as a function of the choice correlation. Results are shown for two values of *p*_CR_, 0.5 and 0.9. **c)** Summary of the relationships between the choice probability CP_*i*_, choice-triggered average CTA_*i*_, and choice correlation CC_*i*_. The picture of how these measures are interconnected is provided by Eqs. 2-6.

Several predictions can be derived from the shape of *h*(*p*_CR_) to experimentally test this threshold-related stimulus dependence of the CP. First, since the influence of *h*(*p*_CR_) is multiplicative, the absolute differences in the CP across different stimulus levels will be higher for cells with a higher CP. Furthermore, the dependence on *h*(*p*_CR_) is fairly flat for a wide range of *p*_CR_ (Fig. 2A), which means that only if including highly informative stimuli in the analysis, leading to *p*_CR_ close to 0 or 1, this dependence will be detectable. For those extreme *p*_CR_ values, CP estimates are less reliable, because for few trials the choice is expected to be inconsistent with the sensory information. This means that, given the number of trials commonly recorded, we expect that only when averaging the CP(*p*_CR_) profile across cells we would be able to detect the modulation of *h*(*p*_CR_). Averaging can also help to average out other cell-specific stimulus dependencies of the choice correlation CC, to isolate the stereotypical factor *h*(*p*_CR_) associated with the threshold effect. Moreover, because *h*(*p*_CR_) multiplies the CC and not the CP, the induced modulation is reversed for negative choice correlations, leading to CPs further decreasing from 0.5 for informative stimulus.

In Fig. 2A we also compare the linear approximation of the CP in terms of the CTA with the exact solution of the extended threshold model (Eq. 14 in Methods). Although derived for weak activity-choice covariations, the approximation is good for a wide range of CP values (see section S1 in the Suppl. Material for further explanations of this goodness). Fig. 2B further compares, as a function of the choice correlation CC, the exact and approximated solutions of the CP. For both the case of an uninformative stimulus (*p*_CR_ = 0.5) and a highly informative stimulus (*p*_CR_ = 0.9) the approximation is accurate for the range of CP values usually found experimentally (0.2 − 0.8). A summary of the overall relation between the CPs, CTAs, and CCs is depicted in Figure 2C.

The multiplicative nature of *h*(*p*_CR_) indicates that the produced stimulus dependence of the activity-choice covariations cannot be eliminated by subtracting an additive stimulus-driven component of the responses, as it is a standard procedure to isolate choice-related signals (Britten et al., 1996; Nienborg and Cumming, 2009; Kang and Maunsell, 2012) when calculating a unique CP combining all stimulus levels. Fortunately, the modulation of the CP due to *h*(*p*_CR_) is relatively small. This implies that the pooling of stimulus-corrected responses is approximately correct and is not expected to introduce major confounds in the estimation of the grand CP when *h*(*p*_CR_) is the only source of activity-choice covariation stimulus dependencies. However, if stimulus dependencies also exist through a stimulus-dependent correlation CC_*i*_(*p*_CR_), a grand CP calculated as an average across stimulus levels will reflect its cell-specific modulation. As we show in section S2 of the Supplementary Material, the grand CP calculated with the pooling of stimulus-corrected responses (Kang and Maunsell, 2012) corresponds to a weighted average of CPs

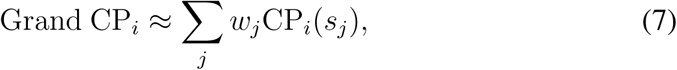

where the weights take the particular form *w*_*j*_ = [*p*(*s*_*j*_|*D* = 1)+*p*(*s*_*j*_|*D* = −1)]/2. This means that reducing the profile of CP_*i*_(*s*) to a single average may introduce potential confounds when grand CP values are compared across cells with different tuning properties, or across recordings from different areas or from different tasks. For example, a high grand CP_*i*_ for a certain cell *i*, compared to others cells, may simply reflect that the sampled stimuli and their weights are such that there is a particularly high CP_*i*_(*s*_*j*\*_) predominant in the weighted average. Conversely, a low grand CP may be due to CP_*i*_(*s*_*j*_) – 0.5 values of opposite sign for different stimuli, which cancel out. In both cases, the high CP for a specific stimulus *s*_*j*\*_ or the asymmetric CP values, are expected to reflect the tuning properties of the cell and its role in the encoding mechanisms, e. g. in the presence of cell-specific stimulus-dependent decision-related feedback (Bondy et al., 2018; Lange and Haefner, 2017). Accordingly, apart from testing the presence of the predicted threshold-induced CP modulation *h*(*p*_CR_), it is also our aim to determine whether our new methods can also identify more cell-specific stimulus dependencies of the activity-choice covariations.

### 2.2 Stimulus dependencies of choice-related signals in the responses of MT cells

We reanalyzed the data from Britten et al. (1996) with new refined methods developed based on the expected properties of the theoretically predicted *h*(*p*_CR_) modulation. We tested for this particular threshold-induced CP stimulus dependence and then more generally characterized the CP(*p*_CR_) patterns found in the data using clustering analysis.

#### 2.2.1 Testing the presence of a threshold-induced CP stimulus dependence

The properties of *h*(*p*_CR_) determine how to examine this CP stimulus dependence. First, because of its multiplicative modulation of the choice correlation, for highly informative stimuli *h*(*p*_CR_) leads to an increase of the CP for cells with positive choice correlation (CPs higher than 0.5) and to a decrease of the CP for cells with negative choice correlation (CPs lower than 0.5). This means that cells with average CP higher and lower than 0.5 should be treated separately or the dependence through *h*(*p*_CR_) would cancel out or reflect the percentage of cells with average CPs higher or lower than 0.5 in the data set. Second, because the effect is strongest for highly informative stimuli, we have to deal with CP estimates with high expected estimation errors, since only in a small percentage of trials will choices be contradictory to the highly informative stimuli. The standard error of 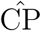 can be approximated as 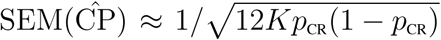 (Bamber, 1975; Hanley and McNeil, 1982, see Methods), where *K* is the number of trials. In the data set the number of trials varies for different stimulus levels, and most frequently *K* = 30 for highly informative stimuli. In that case, for *p*_CR_ = 0.9, only 3 trials for choice *D* = −1 are expected, and 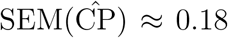. As can be seen from Fig. 2, this error surpasses the order of magnitude of the CP modulations expected from *h*(*p*_CR_). Therefore, we need to average the CP(*p*_CR_) profiles across cells not only to balance out other potential cell-specific CP stimulus dependencies, but also to reduce the standard error. This average should take into account a third property of the *h*(*p*_CR_) modulation, namely that its effect is relative to the value of CP(*p*_CR_ = 0.5). Therefore, the average across cells should only include cells for which a CP(*p*_CR_) can be calculated for each value of *p*_CR_ in the range under consideration. Otherwise, the shape across *p*_CR_ values of the CP average across cells would mostly reflect the changes in the distribution of CP(*p*_CR_ = 0.5) values among the particular subset of cells contributing to the CP average for each particular *p*_CR_ value.

Given these points, we analyzed the CP(*p*_CR_) dependencies as follows. First, for each cell and each stimulus coherence level we calculated a CP estimate if at least 4 trials were available for each decision. We then binned the range of *p*_CR_ into five bins, [0 − 0.3, 0.3 − (0.5 − *ε*), (0.5 − *ε*) − (0.5 + *ε*), − (0.5 + *ε*) − 0.7, 0.7 −1]. The value of *ε* was chosen such that only trials with the uninformative (zero coherence) stimulus were comprised in the central bin. Wider bins were selected at the extremes of the range to compensate for higher 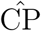 standard errors. The following results are robust to the selection of the minimum number of trials and the binning intervals. CPs estimated at different coherence levels were assigned to the bins according to the psychometric function, and a weighted average CP per bin was calculated. The weights were assigned as inversely proportional to the standard error of the estimates. This type of weighted average is used throughout our analysis when combining CP values across stimulus levels or cells (see Methods for details). This procedure should produce for each cell a CP(*p*_CR_) curve with five dots, one per bin. However, for many cells in the data set this curve could not be completed because the requirement on a minimum number of trials precluded from estimating a CP value for some of the bins. We obtained a complete curve for *N* = 107 cells. If not stated otherwise, subsequent analyses focus on this subset of cells. Average CP(*p*_CR_) profiles across these cells were calculated separately for cells with an average CP across *p*_CR_ higher or lower than 0.5.

Fig. 3A shows the averaged CP(*p*_CR_) profiles. To assess the statistical significance of the observed CP dependencies on *p*_CR_ we developed a new method to construct surrogate data. These surrogates allow testing whether a pattern consistent with the predicted CP increase for informative stimuli could appear under the null hypothesis that the CP has a constant value independent of *p*_CR_ (see Methods for details). For the cells with average CP higher than 0.5, we found that the modulation of the CP was significant, in agreement with the model. For cells with average CP lower than 0.5 the modulation was not significant. This difference could be explained by the following reasons. First, from a statistical point of view, the fact that from the *N* = 107 cells included in this analysis 74 had a CP higher than 0.5 and 33 lower, means that the estimation error of the average is bigger for the average of the cells with CP < 0.5. Second, if the modulation is multiplicative as predicted by *h*(*p*_CR_), its impact is expected to be smaller when the magnitude of the activity-choice covariations is smaller. Indeed, CP values on average are closer to 0.5 for the cells with CP < 0.5 (see Fig. 3A, consistently with the results in Figure 5 of Britten et al. (1996)). We further confirmed the robustness of the CP modulation in a wider set of cells. For this purpose, we repeated the analysis forming subsets separately including cells with a calculable CP for the three bins with *p*_CR_ lower or equal 0.5, and the three with *p*_CR_ higher or equal than 0.5. Also in this case the observed CP(*p*_CR_) pattern was robust for cells with average CP higher than 0.5 (Fig. 3B, *N* = 171).

**Figure 3:**
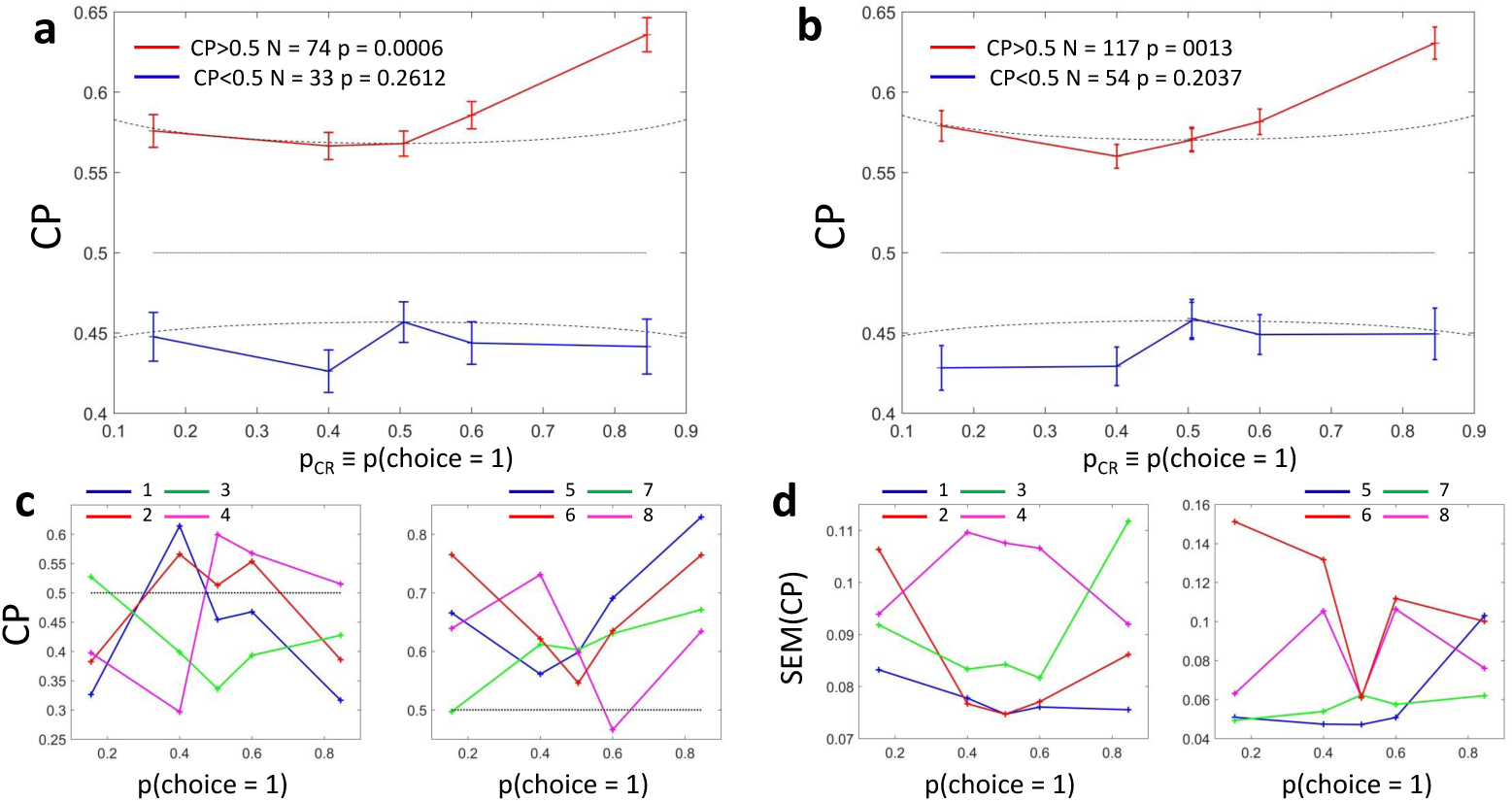
Choice probability as a function of the ratio of choices for MT cells during a random dots discrimination task (Britten et al., 1996). **a)** Average CP for five bins spanning the range of *p*_CR_ ≡*p*(*D* = 1) (see main text). *N* = 107 cells were preselected based on the criterion that CP values could be estimated for all bins. The average was calculated separately for cells with average CP higher or lower than 0.5. Dotted lines reflect the relationship predicted by the factor *h*(*p*_CR_) (Eq. 5). Significance of the stimulus dependencies was evaluated against the null hypothesis of a constant CP value using surrogate data (see Methods). **b)** Same analysis but with a less demanding criterion of inclusion that separately averages across cells with CP values available for bins corresponding to *p*_CR_ values lower or higher than *p*_CR_ = 0.5. **c)** CP(*p*_CR_) profile for four example cells with average CP lower and higher than 0.5, respectively. **d)** Standard error of the estimated CP for the example cells as a function of *p*_CR_.

**Figure 4:**
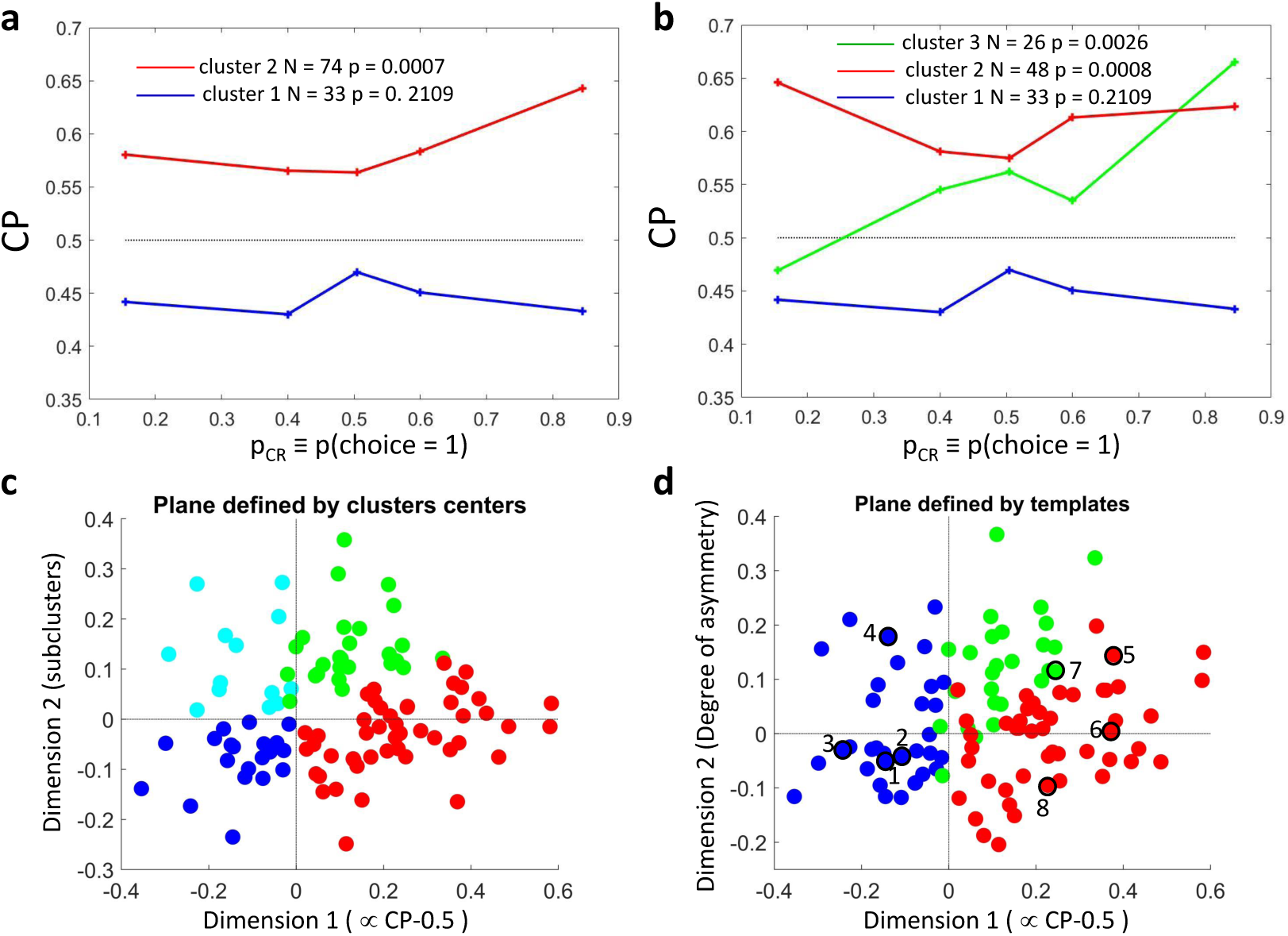
Symmetric and asymmetric dependencies of choice probability as a function of *p*_CR_. **a-b**) CP as a function of *p*_CR_ for clusters of the MT cells determined by *k*-means clustering. Each CP(*p*_CR_) profile corresponds to the center of a cluster. Significance of the modulation was quantified as in Figure 3. **a**) Two clusters (*N*_*c*_ = 2) for all cells. **b**) Further subclustering of cells with average CP > 0.5 into two subclusters. **c-d**) Representation of the CP(*p*_CR_) profiles in a two-dimensional space spanned by the cluster means. The horizontal axis is defined by clusters 1 and 2 and closely aligned with CP − 0.5. **c**) Vertical axis is defined as perpendicular to horizontal axis in the plane defined by the subcluster means. Colors correspond to the clusters of panel b, with blue & cyan further indicating subclusters of cells with average CP < 0.5 (see Fig. 7A in Supplementary Material). **d**) Space defined by projection onto two templates: a constant (x-axis) representing CP magnitude, and identity (y-axis) representing CP asymmetry. Colors correspond to the clusters of panel b and numbers to the examples of Figure 3C.

**Figure 5:**
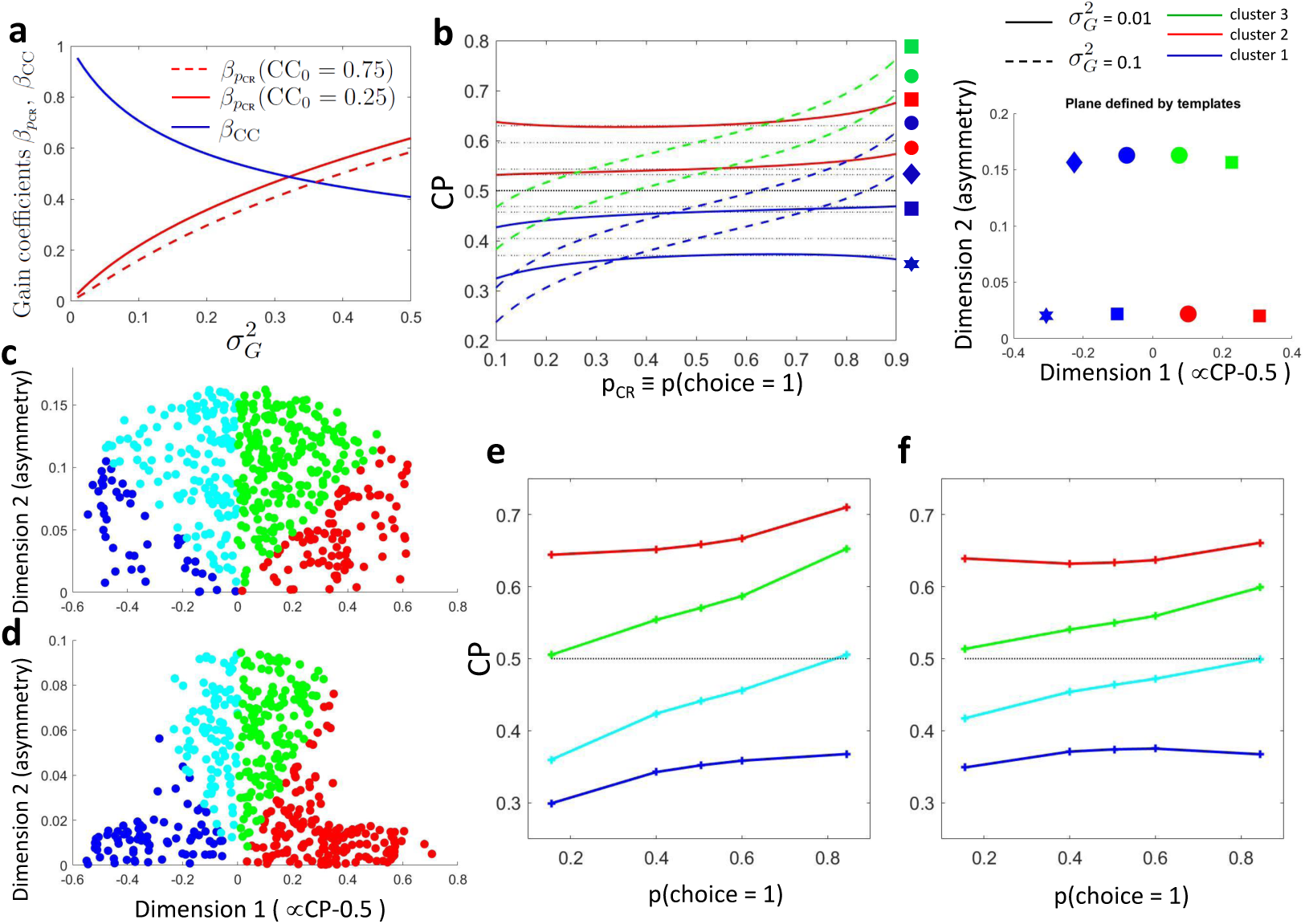
Modeling the influence of neuronal gain modulation on CP(*p*_CR_) profiles. **a**) Dependence of gain coefficients *β*_*CC*_ and *β*_*p*CR_ (Eqs. 8 and S20) on the strength of the gain fluctuations, 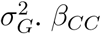 determines their effect on the choice correlation CC_*i*_(*s*_0_) for the uninformative stimulus *s*_0_. *β*_*p*CR_ determines the degree of asymmetry of the choice correlation dependence on the *p*_CR_. **b**) CP(*p*_CR_) profiles for different combinations of CC_*i*0_(*s*_0_) and 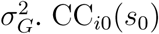 is the choice correlation that would be obtained for *s*_0_ with no gain fluctuations. We display CP(*p*_CR_) for four values of CC_*i*0_(*s*_0_) (lines vertically separated) and two values of 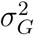 (solid vs dashed). Each line represented by a symbol corresponds to a point in the two-dimensional space defined by the symmetric and asymmetric templates introduced in Fig. 4D. **c**) CP(*p*_CR_) profiles, represented in the same 2-D space, generated with a uniform sampling of CC_*i*0_(*s*_0_) consistent with the observed average CPs of the MT cells, and with a uniform sampling of the gain 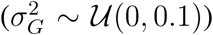. **d**) Analogous to c, but with a nonuniform distribution in the 2-D space, reflecting structure in the covariation of CC_*i*0_(*s*_0_) and 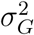. **e**-**f**) CP(*p*_CR_) profiles corresponding to the clusters centers obtained when sampling the space according to panels c and d, respectively.

These results provide evidence supporting an increase of the CP for informative stimuli. However, the dependence found for cells with CP > 0.5 only partially corresponds to the shape predicted by *h*(*p*_CR_). In particular, the CP increase appears to be higher for *p*_CR_ > 0.5. The finding of this asymmetry is consistent with results reported in Britten et al. (1996), who found a significant but modest effect of coherence direction on the CP (see their Figure 3). By experimental design, the direction of the dots corresponding to choice *D* = 1 was tuned for each cell separately to coincide with their most responsive direction. This means that this asymmetry indicates that CPs tend to increase more when the stimulus provides evidence for the direction eliciting a higher response. However, Britten et al. (1996) found no significant relation between the magnitude of the firing rate and the CP (see their Figure 3), and we confirmed this lack of relation specifically for the subset of *N* = 107 cells (result not shown). This eliminates the possibility that higher CPs for high *p*_CR_ > 0.5 values are due only to the higher responses, and suggests a richer underlying structure of CP(*p*_CR_) patterns. Given that our refined methods are designed to better characterize the within-cell CP(*p*_CR_) profiles, we next combined them with nonparametric clustering to characterize any structure beyond the specific threshold-induced predicted modulation.

#### 2.2.2 Characterizing the patterns of CP stimulus dependencies with cluster analysis

Beyond the evidence obtained from the average CPs of the presence of a CP dependence on the stimulus level, screening CP(*p*_CR_) profiles for individual cells shows substantial heterogeneity across cells (Fig. 3C). Across cells, the modulation predicted by *h*(*p*_CR_) did not account for more variance than a model assuming that the CP was constant. In particular, the coefficient of determination *R*^2^, quantifying the difference in accounted variance across *p*_CR_ values between the model using *h*(*p*_CR_) and a model assuming a constant CP value, was on average −0.07 and −0.008 for cells with average CP lower and higher than 0.5, respectively. The CP(*p*_CR_) profiles of individual cells vary with respect to *h*(*p*_CR_) (Fig. 3C). However, these low *R*^2^ values are not incompatible with the significant modulation of the average shown in Fig. 3A, B, since the variability across cells is averaged out at the population level. This diversity across cells suggests that together with a stereotypical modulation *h*(*p*_CR_), other cell-specific sources of stimulus dependence may exist. This cell-specific dependencies could be caused, for example, by changes in the strength of feedback choice-related signals depending on the estimate of the stimulus (Bondy et al., 2018; Lange and Haefner, 2017). However, given the high 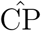 standard errors for the single cells (Fig. 3D), a substantial part of this variability may also not reflect any underlying computational mechanisms.

To further assess the existence of CP stimulus dependencies, we carried out unsupervised *k*-mean clustering (Bishop, 2006) to examine the patterns of CP(*p*_CR_) without *a priori* assumptions about a modulation *h*(*p*_CR_) associated with the threshold effect. Clustering was performed considering the profile of CP(*p*_CR_) −0.5 for each cell as a vector in a 5-dimensional space, where 5 is the number of bins, as described above. To consider both the shape and sign of the modulation, distances were calculated with the cosine distance (one minus the cosine of the angle between the vectors), and clustering was repeated for a range of pre-specified number of clusters. With two clusters the distinction between cells with CP higher or lower than 0.5 is naturally recovered (Fig. 4A). Significance was again assessed constructing surrogate CP(*p*_CR_) profiles and repeating the clustering analysis on these surrogates. Again a significant dependence of the CP on *p*_CR_ was found only for the cluster associated with CP higher than 0.5.

To further separate CP(*p*_CR_) patterns, we iterated the clustering procedure to divide each of the two clusters into subclusters. In the following, results are shown for the case in which this procedure is applied separately to cells with average CP higher and lower than 0.5, given that only for the former the modulation is found significant. Average CP(*p*_CR_) profiles for the two subclusters of cells with CP > 0.5 are shown in Fig. 4B. For both subclusters the CP(*p*_CR_) dependence is significant. The larger cluster has a more symmetric shape of dependence on *p*_CR_, with an increase of CP in both directions when the stimulus is informative. Conversely, for the smaller cluster the dependence is asymmetric, with an increase when the stimulus information content is consistent with the preferred direction of the cells and a decrease in the opposite direction.

Introducing a second cluster allows us to represent each vector of CP(*p*_CR_) in a two dimensional space (Fig. 4C). The horizontal axis corresponds to the separation between the two initial clusters, and is closely aligned to the departure of the average CP from 0.5. The vertical axis is defined by the vectors corresponding to the centers of the two subclusters and is determined separately for the cells with average CP higher and lower than 0.5 (see Methods for details). For cells with CP > 0.5 this second axis can be associated with the degree to which the CP(*p*_CR_) dependence is symmetric or asymmetric with respect to *p*_CR_ = 0.5. Cells for which the CP increases consistently with its preferred coherence lie on the positive plane. To further support this interpretation of the axis, we repeated the clustering procedure replacing the nonparametric *k*-mean procedure with a parametric procedure selecting a priori a symmetric and an asymmetric template for the subclusters. As can be seen from the comparison of Fig. 4C-D, the space obtained with template clustering reproduces well the representation in the space obtained nonparametrically. For the cells with CP < 0.5, the separation between the two subclusters has a less clear interpretation, consistently with the lack of significance of the CP(*p*_CR_) dependence (see Fig. 7A in Supplementary Materials).

Similar results are obtained when increasing the number of clusters keeping all cells together, instead of separating cells with average CP higher and lower than 0.5. Introducing a third cluster for all cells leaves almost unaltered the cluster of cells with CP lower than 0.5 (Fig. 7B in Supplementary Material). The cluster of cells with CP higher than 0.5 splits in two subclusters analogous to the ones found from cells with CP higher than 0.5 alone. The distinction between cells with more symmetric and asymmetric CP(*p*_CR_) dependencies is robust to the selection of a larger number of clusters, i.e. clusters with this type of dependencies remain large when allowing for the discrimination of more patterns (Fig 7C in Supplementary Materials).

This clustering analysis confirms the presence of stimulus-dependent choice-related signals, and further indicates that these dependencies have a structure beyond the stereotypical dependence *h*(*p*_CR_). While the stimulus dependence *h*(*p*_CR_) is generic for the threshold model, other CP stimulus dependencies through the choice correlation CC will depend on the source of choice-related signals. The structure of CP stimulus dependencies could reflect, for example, the structure of noise correlations or of feedback projections. A more complete characterization of the structure of these stimulus dependencies would require a data set systematically tiling the space of receptive fields and is beyond the scope of this work. Alternatively, we here show that some main features of the patterns observed in Fig. 4 can be captured with still a generic threshold model without a comprehensive specification of the properties of connectivity and noise correlations.

In particular, we focus on two main features of these CP(*p*_CR_) dependencies observed for the MT cells. First, the existence of asymmetric CP stimulus dependencies. Second, that the average magnitude of the CP and the degree of asymmetry of CP(*p*_CR_) negatively covary, as seen when comparing the more symmetric (red) and asymmetric (green) profiles in Fig. 4B. To capture these two features with a simple model, we studied a linear threshold model (Shadlen et al., 1996; Haefner et al., 2013) in which the sensory neurons are subjected to global fluctuations in their excitability (see Methods for details). Goris et al. (2014) showed that this source of variability can account for a substantial amount of the variance of individual cells in visual sensory areas. In particular, they estimated that a 75% of the variability in the responses in monkeys MT cells when presented with drifting gratings could be explained by gain fluctuations. We applied their method to estimate the strength of gain fluctuations in the data set of Britten et al. (1996) and found that approximately half of the variance could be attributed to gain fluctuations. It is thus pertinent to examine how these fluctuations can affect activity-choice covariations.

We now show that a model accounting for gain fluctuations produces asymmetric CP(*p*_CR_) patterns and a link between the magnitude of the CP and the shape of CP(*p*_CR_) profiles. Fluctuations introduce a component of the noise covariance matrix Σ proportional to the tuning curves (∝**f**(*s*)**f**^*T*^(*s*), see Methods). This component renders the covariance matrix stimulus dependent, i. e. Σ(*s*), which alters the activity-choice covariation cov(*r*_*i*_, *d*) and the variance var *r*_*i*_ of the responses for each stimulus level. This stimulus dependence is inherited by the choice correlation (Eq. 4) and hence by the CP. The strength 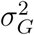 of the gain fluctuations determines the strength of this cell-specific stimulus dependence. Furthermore, the strength of the gain also affects the magnitude of the CP, by adding variability to the responses unrelated to the choice. Therefore, the strength of the gain introduces a covariation between the CP magnitude and its stimulus dependencies. In more detail, we analyzed a purely feedforward threshold model with optimal read-out weights (Haefner et al., 2013; Pitkow et al., 2015) and considered the effect of gain fluctuations (see Methods and section S4 in the Suppl. Material). For this model, we obtained the following stimulus dependencies:

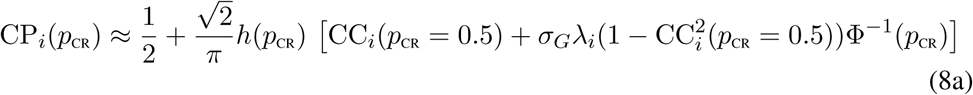

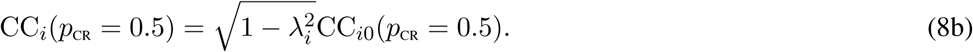

Here 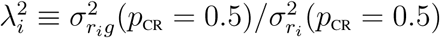 is the portion of the responses variance of cell *i* caused by the gain fluctuations. The variance due to gain fluctuations is 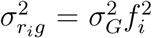(Eq.22). CC_*i*0_(*p*_CR_ = 0.5) is the choice correlation that cell *i* would have if there were no gain fluctuations 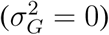.

The stimulus dependence of CP(*p*_CR_) appears, like in the case without gain fluctuations, through the factor *h*(*p*_CR_), common to all cells, which is symmetric. Moreover, now an asymmetric dependence is also determined by Φ^−1^(*p*_CR_), which maps the *p*_CR_ to the associated stimulus *s*. The coefficient of this factor, 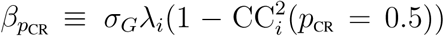, is cell-specific. The choice correlation CC_*i*_(*p*_CR_ = 0.5) also depends on the strength of the gain. The coefficient 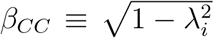 modulates CC_*i*0_(*p*_CR_ = 0.5). The covariation between the magnitude of the CP and the degree of asymmetry in CP(*p*_CR_) occurs as follows: when 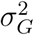 increases, 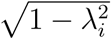 decreases, and thus CC_*i*_(*p*_CR_ = 0.5) decreases. Conversely, the increase of 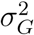 increases both *λ*_*i*_ and 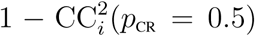, and thus the coefficient *β*_*p*CR_ increases, making the dependence of CP(*p*_CR_) on Φ^−1^(*p*_CR_) stronger. Therefore, cells with CP_*i*_(*p*_CR_ = 0.5) closer to 0.5 have a stronger asymmetric dependence on *p*_CR_. Furthermore, a CP_*i*_(*p*_CR_ = 0.5) closer to 0.5 leads to a smaller effect of the multiplicative symmetric modulation *h*(*p*_CR_), further contributing to the negative covariation between the magnitude and degree of asymmetry.

The covariation of the coefficients *β*_*p*CR_ and *β*_*CC*_ that modulate the strength of the CP(*p*_CR_) dependence and the magnitude of CP_*i*_(*p*_CR_ = 0.5), respectively, is shown in Fig. 5A. To determine 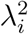 only in terms of the strength of the gain we fixed the rate *f*_*i*_(*p*_CR_ = 0.5) = 10 spike/s and considered the variance not associated with the gain to be poissonian 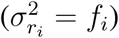, so that 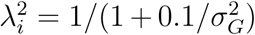. The range 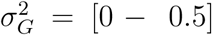 corresponds to 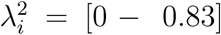. Fig. 5B shows CPs as a function of *p*_CR_ for combinations of four values of CC_*i*0_(*p*_CR_ = 0.5) and two values of 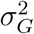. The other parameters of the cell responses are kept constant as in Fig. 5A, so that 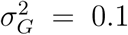 corresponds to 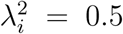 and 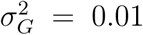 to 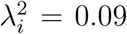. The representation of these CP(*p*_CR_) profiles in the two dimensional space defined by the same symmetric and asymmetric templates used in Fig. 4D is also displayed. The model qualitatively reproduces the presence of symmetric and asymmetric clusters, with a negative correlation between the magnitude and the degree of asymmetry of the CP(*p*_CR_) profile. Fig. 5C-F further illustrate how combinations of different CC_*i*0_(*p*_CR_ = 0.5) and 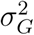 populate the 2-D space of CP(*p*_CR_) profiles. CP(*p*_CR_) profiles were simulated randomly sampling the average CP values from the ones observed for the MT cells. For Fig. 5C,E, the fluctuation gains were uniformly sampled from the interval 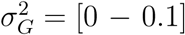, corresponding to 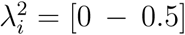. For Fig. 5D,F, the 2-D space was not evenly sampled, simulating a further dependence between CC_*i*0_(*p*_CR_ = 0.5) and 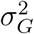 values which determines the exact balance between the symmetric and asymmetric dependencies observed in the average profiles associated with each cluster.

This simple model provides a plausible explanation for two main features observed in the CP(*p*_CR_) patterns of the MT cells, namely the existence of asymmetric CP(*p*_CR_) dependencies and of a negative correlation with the CP magnitude. The model does not require extra specifications of the properties of the tuning functions or the connectivity and noise correlations structure. However, modeling other aspects such as the fact that for the MT cells the CP(*p*_CR_) dependencies observed are weaker for cells with average CP < 0.5 would demand further adding a specific structure to the space of CP(*p*_CR_) profiles, namely such that *λ*_*i*_ was lower for the cells with CP < 0.5. Heterogeneity in the proportion *λ*_*i*_ of variance explained by the gain is expected to be associated with the structure of connectivity and noise correlations. In particular, while in the model we assumed that the strength of the gain fluctuations was common to the whole population of sensory neurons, the effective cell-specific impact of the fluctuations in the CP is expected to depend on that structure, since the CP of a cell also depends on fluctuations in the excitability of those cells with highly correlated activity, as determined by the noise correlations (Haefner et al., 2013). Further characterizing the CP(*p*_CR_) patterns of MT neurons in terms of the structure of the connectivity and noise correlations would require data from simultaneous recordings systematically tiling the space of the cells response tuning functions.

### 2.3 Modeling stimulus-dependent choice-related signals with GLMs

The analysis of CP stimulus dependencies in the data from Britten et al. (1996) indicates that the stimulus modulates choice probabilities. Next we examine how accounting for these stimulus dependencies can improve statistical models of activity-choice covariations.

In particular, we study how stimulus-dependent choice-related signals affect the fit of generalized linear models (GLMs) of neural responses (Truccolo et al., 2005). Although CPs have been a traditional way to quantify the covariation of neural responses with the choice, GLMs provide a more versatile framework in which these covariations can be modeled jointly with the dependence on other explanatory variables, such as the external stimulus, response memory, or interactions across neurons (Park et al., 2014; Runyan et al., 2017). Typically, GLMs are constructed in a modular way, such that each of these explanatory variables contributes with a multiplicative factor that modulates the firing rate of a Poisson process. With this type of implementation, the choice modulates the rate as a binary gain factor, with a different gain for each of the two choices. Its multiplicative nature already introduces some covariation between the impact of the choice value on the rate and the one of the value of the other explanatory variables. However, using a single coefficient (or kernel) to model the effect of the choice on the neural responses may be insufficient if choice-related signals are stimulus dependent in more complex ways, as suggested by the CP stimulus dependencies that we observed.

To evaluate how accounting for stimulus dependencies of choice-related signals could improve the fitting of GLMs to the MT cells responses, we compared the likelihood of three types of nested models. In the first type, the rate in each trial is predicted only based on the external stimulus level. In a second type, the effect of the choice is incorporated, but it contributes with a single predictor. In a third type, multiple choice-related terms are introduced, with an indicator function that indicates which is the factor that is active for a certain subset of stimulus levels. In more detail, the full model has the following form for the mean rate *μ*(*r*_*i*_) of the responses of cell *i*:

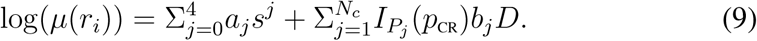

The terms 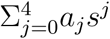 model the stimulus influence with a fourth order polynomial, and are the only constituents of the first type of model. The choice dependence is modeled by 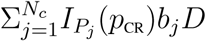, with *N*_*c*_ ∈ [1, 3] parameters. 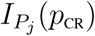 is an indicator function which equals one if the *p*_CR_ value belongs to the subset *P*_*j*_ associated with the choice parameter *b*_*j*_, and is zero otherwise. For the second type of models *N*_*c*_ = 1 and hence the choice affects the predicted responses equally for all stimulus levels. For the third type of model, when fitting multiple choice parameters, we determined the subsets of stimulus levels associated with each of those parameters using the CP(*p*_CR_) profiles for a first characterization of the stimulus dependencies (see Methods for further details). Like for the CP analysis, for each cell we determined which coherence values could be included in the analysis given a criterion requiring a minimum number of trials for each choice (at least 4). To avoid overfitting, we considered only models with *N*_*c*_ = 1, 2, 3. When having multiple parameters instead of a single coefficient *b*, the vector *b*(*p*_CR_) with components *b*_*j*_, *j* = 1, …, *N*_*c*_ reflects stimulus dependencies analogously to the CP(*p*_CR_) profile.

To compare the models, we separated the data into training and testing sets, and calculated the average likelihood for each type of model using cross-validation to account for overfitting. To quantify the increase in predictability when adding the choice as a predictor we defined the likelihood relative increase (LRI) as [*L*(*choice, stimulus*) − *L*(*stimulus*)]/[*L*(*stimulus*) − *L*_0_], where *L*_0_ is the likelihood of a model assuming a constant rate across all stimulus levels and no choice modulation. That is, the LRI quantifies the relative increase of further adding the choice as a predictor relative to the increase of previously adding the stimulus as a predictor. This measure can be interpreted as an indicator of the relative influence of the choice and the sensory input in the neural responses, hence expected to increase from sensory to higher perceptual areas. Fig. 6A compares the LRI values for models with a single choice parameter versus models that allow for multiple choice parameters. LRIs are predominantly higher when allowing for multiple parameters. We summarized these results in two ways: Fig. 6B shows the proportion of cells in each cluster for which the LRI was higher than a ten per cent and Fig. 6C shows the average LRI values. Both reflect the increase in LRIs when using multiple choice parameters.

**Figure 6:**
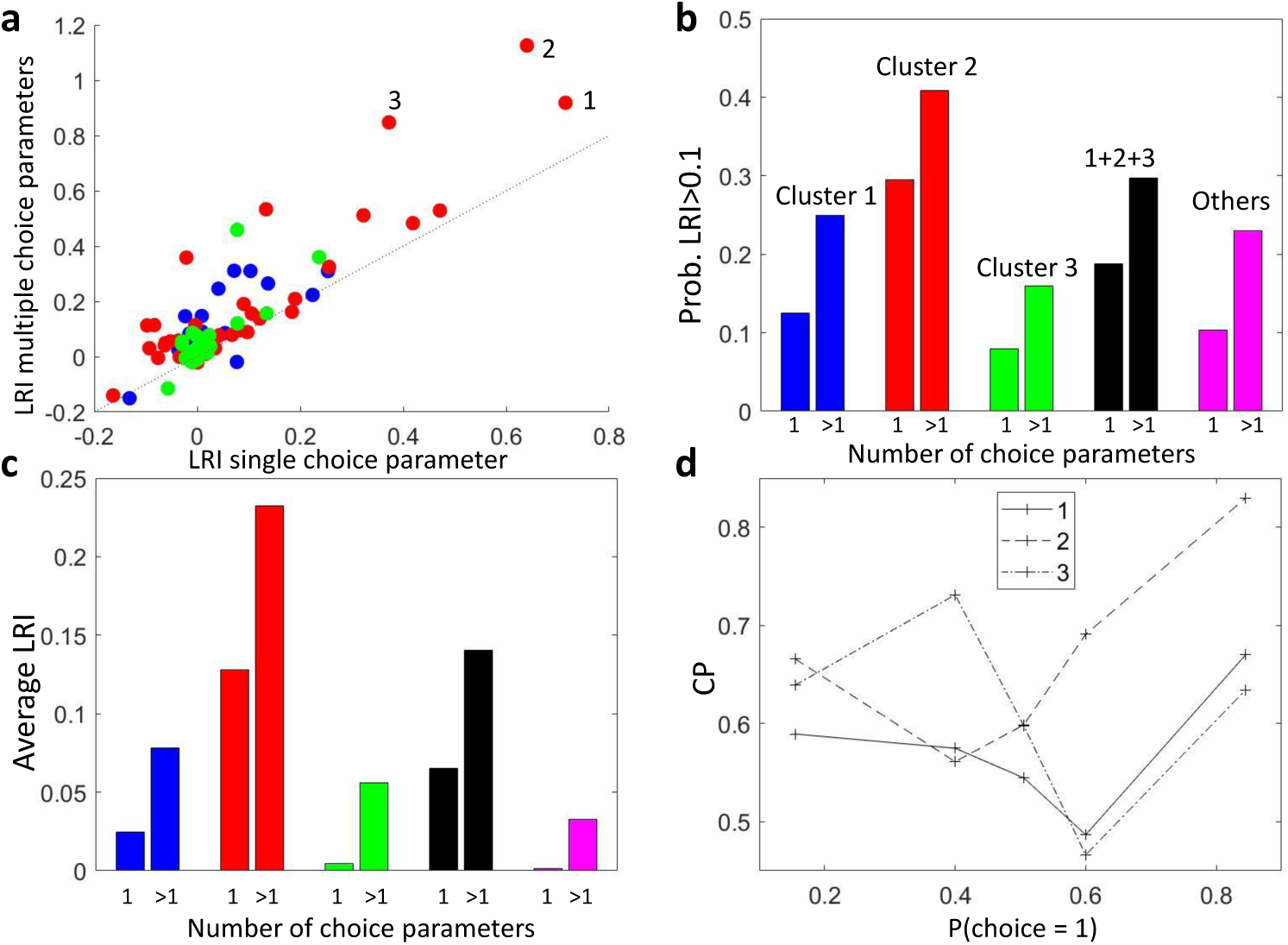
Modeling stimulus-dependent choice-related signals with GLMs. **a**) Raster plot of the cross-validated likelihood relative increase (LRI) of models with a single choice parameter versus models with multiple choice parameters associated with different stimulus levels. **b**) Proportion of cells with LRI> 0.1 for models with single and multiple choice parameters, grouped by the clusters as in Fig. 4B. Cells not included in the set of 107 cells for which a CP value could be estimated for each bin of *p*_CR_ are labeled as ‘Others’. **c**) Average LRI values, grouped as in b. **d**) CP(*p*_CR_) profiles of the three cells with the highest LRI in the models with multiple parameters, as numbered in panel a.

There are two reasons why accounting for stimulus-choice interaction terms in GLMs can help to characterize the activity-choice covariations. First, the patterns of *b*(*p*_CR_) parameters, similarly to CP(*p*_CR_) patterns, may be expected to have a structure associated with specific cell properties and their role in the perceptual decision-making process. Second, even when the characterization of activity-choice covariations is reduced to a single measure of strength, the use of GLMs containing interaction terms will reassess the comparison across cells of how stimulus driven versus choice driven different cells are, since these terms will improve the fitting to a different degree for different cells, as a function of how stimulus dependent the choice-related signals are.

To illustrate this, we can compare how adding the interaction terms affects the ranking of the three cells with highest LRI when using multiple choice parameters (Fig. 6A), which CP(*p*_CR_) profiles are shown in Fig. 6D. Although these cells also have high LRI values with a single parameter, the rank of cell 1 and 2 flips because of the higher CP(*p*_CR_) modulation of cell 2. Similarly, while the LRI with multiple parameters for cell 1 and 3 are close, the LRI of cell 3 is substantially lower with a single parameter, indicating that its pattern of stimulus dependence is less well captured by a single parameter. The degree to which a model with interaction terms improves the predictability will depend on the shape of the CP(*p*_CR_) patterns, which themselves are expected to vary across areas or across cells with different tuning properties. For example, we see in Fig. 6C that for the cluster with an asymmetric CP(*p*_CR_) profile (cluster 3), the average LRI with only one choice parameter suggests that this type of cells are not choice driven. This can be understood because for the cells in this cluster the sign of the choice influence on the rate is more stimulus dependent, and a single choice parameter gives a worse prediction for choice influences of opposite sign at different stimulus levels. Overall, accounting for stimulus-choice interactions in GLMs is expected to provide a more accurate comparison across neurons of the relative importance of stimulus and choice influences in the responses.

## 3 Discussion

Our work makes several contributions to the understanding of how choice and stimulus signals in neural activity are coupled. The first is that we analyzed a general model of perceptual decision-making to analytically derive how the relationship between sensory responses and choice depends on stimulus strength, regardless of whether this relationship is due to feedforward or feedback choice signals, when the choice is estimated from sensory responses through a threshold mechanism. Second, we used the model insights to design a new, more sensitive, methodology to measure the dependence of choice probabilities (CPs) on stimulus strength. Third, we used this methodology to test our predictions using the classic dataset by Britten et al. (1996). Interestingly, we found a richer structure in how CPs in MT neurons depend on stimulus strength than expected. In addition to a symmetric dependency predicted by the model, we found an asymmetric dependency which we could explain by incorporating previously observed gain fluctuations (Goris et al., 2014) into our model. Fourth, we showed that generalized linear models (GLMs) that account for stimulus-choice interactions better explain sensory responses in MT and allow for a more accurate characterization of how stimulus-driven and how choice-driven a cell’s response is.

Previous work has demonstrated that solving analytically models of perceptual decision-making leads to important new insights on the interpretation of the relationship between neural activity and choice (Bogacz et al., 2006; Gold and Shadlen, 2007; Haefner et al., 2013). In particular, in previous work Haefner et al. (2013) computed analytical CPs in a decision-making model in which sensory responses give rise to choices through a decision threshold operating upon a feed-forward linear read-out, when the two choices occur with equal probability. This earlier work has been instrumental in better interpreting CPs, and in showing how experimentally measured CPs relate to the read-out weights that the neurons have in the stimulus internal decoder. Here we provided a general analytical solution of CPs in a more general model, valid in the presence of both feedforward and feedback choice signals. Moreover, our solution is valid when the stimulus is informative and produces any ratio of choices, that is, when the rate of each choice over trials is different. Accordingly, we derived the analytical dependency of CP on the probability of one of the choices (*p*_CR_ ≡ *p*(choice = 1)), which determines the ratio of choices and mediates the dependency of the CP on the stimulus strength. Our model is therefore directly applicable to both discrimination and detection tasks, for any stimulus strength that elicits both choices. As we illustrated, these advances in the analytical solution of the decision-threshold model proved to be very helpful to detect and interpret the stimulus dependency of choice-related signals in neural activity.

One immediate practical outcome of the analytical solution of our model was that it allowed us to understand some difficulties that have clouded previous attempts to find stimulus dependencies of CPs in real neural data. Our model showed that the predicted direction of the dependence of CP on *p*_CR_ is different for neurons with CP larger or smaller than chance (that is, neurons more responsive for opposite choices). This opposite modulation would greatly reduce the magnitude of the overall dependence of the CP on stimulus strength when averaging over all neurons, as done in previous analyses (Britten et al., 1996). Furthermore, the magnitude of the dependence of CPs on *p*_CR_ also depends on how far is the CP from chance level. This implies that neuron-specific dependencies should be characterized for each cell individually relative to the CP obtained with the uninformative stimulus. Only neurons for which a full individual CP profile can be estimated should be combined to determine stimulus dependencies at the population level, or otherwise the overall profile of dependence will be dominated by variability associated with the different subsets of neurons contributing to the estimate at each stimulus level. Informed by these insights we could derive more refined methods with higher statistical power to characterize the within-cell dependencies of choice-related signals on stimulus strength^2^. The application of these methods to the classic neural data from MT neurons during a perceptual decision-making task of Britten et al. (1996) allowed us to find stimulus dependencies of CPs, while the simpler analysis pooling all neurons failed to find a significant effect.

Our understanding of how CP-stimulus dependencies may arise within the decision-making process, and the resulting new methods to measure these dependencies on data, should contribute to refine how CPs in neural populations are measured and interpreted. Traditional analyses take grand averages across stimulus levels and obtain a distribution of CPs, with a single CP value per neuron, within a certain area or neural population. Areas or populations are then ranked in terms of these grand-averaged CP values, to determine decision-making areas as those areas which present a higher association between neural activity and choice. In these calculations, it is a standard procedure to assume no effect of the stimulus signal on the activity-choice covariation (e.g. Nienborg and Cumming, 2009; Pitkow et al., 2015; Wimmer et al., 2015; Smolyanskaya et al., 2015) and to calculate a single grand CP (Britten et al., 1996), combining trials from all stimulus levels. This assumption has also been motivated by practical reasons, because activity-choice covariations for single sensory neurons are small and pooling trials across stimulus levels helps to obtain a better estimate of the average CP. The grand CP is calculated directly as a weighted average of the CPs estimated for each stimulus level, or, as most commonly, pooling the responses from trials of all stimulus levels, after subtracting the stimulus-related component (Kang and Maunsell, 2012). We have used our theoretical CP analysis to show that the latter procedure in fact corresponds to a specific type of weighted average, with the weights determined by which stimuli are most frequent for each choice. Either way, in the presence of a non-constant CP(*p*_CR_) profile, the weighted average may introduce confounds in the assessment of the average strength. For example, CP(*p*_CR_) patterns with different sign for different *p*_CR_ values will result in lower average CP values. The comparison of the average CP of a cell across tasks may mostly reflect changes in the sampling of stimulus levels. Similarly, if the structure of CP(*p*_CR_) patterns is related to the tuning properties, the comparison of the average CP across cells with different tuning properties will mostly depend in the sampling of stimulus levels. This limitation is not specific to average CP values, and applies to other measures that consider choice-related and stimulus-driven components of the response as separable, such as partial correlations (e. g. Zaidel et al., 2017). Our work instead indicates that the shape of the CP(*p*_CR_) patterns cannot be summarized in the average, and this shape may be more informative about the role of the activity-choice covariations, when comparing across cells with different tuning properties, cells from different areas, or across task variations (e.g. Romo and Salinas, 2003; Nienborg and Cumming, 2006; Nienborg et al., 2012; Krug et al., 2016; Sanayei et al., 2018; Shushruth et al., 2018; Jasper et al., 2019; Steinmetz et al., 2019). Our new methods allow individuating and quantifying these CP patterns and hence a better characterization of the covariations between neural activity and choice across neural populations.

Our work has also implications for improving generalized linear models of neural activity, which are commonly used to describe neural responses in the presence of many explanatory variables that could predict the neuron’s firing rate, such as the external stimulus, motor variables, autocorrelations or refractory periods, and the interaction with other neurons (Truccolo et al., 2005; Park et al., 2014; Runyan et al., 2017). While usually the stimulus and the choice are treated as separate explanatory variables, we extended the GLMs to include explicit interactions between choice and stimulus. We showed that, consistently with the finding of non-constant CP(*p*_CR_) patterns, these models improved the goodness of fit for the responses of the MT cells. Importantly, properly accounting for stimulus-choice interaction terms affected the quantification of how stimulus-driven or choice-driven different cells are, quantified as the increased in predictive power when further adding the choice as a predictor after the stimulus. This progress offers a simple way to better evaluate which sets of neurons (e. g. across areas, or cell types) have a higher association with the behavioral choice or the stimulus. Our refined GLMs with multiple choice parameters associated with subsets of stimulus levels allow characterizing the patterns in the vector of choice parameters analogously to CP(*p*_CR_) patterns. Furthermore, our approach can be extended straight-forwardly to GLMs that model stimulus dependencies of choice-related signals across the time-course of the trials, using more complex kernels to account for the effect of experimental covariates on the neural responses (Park et al., 2014). This implies that GLMs with stimulus-choice interaction terms can also be applied to experimental settings with multiple sensory cues presented at different times (e.g. Romo and Salinas, 2003; Sanayei et al., 2018) or a continuous time-dependent stimulus (Nienborg and Cumming, 2009). Similarly, the interaction terms may also help to model the influence of choice history in the processing of sensory evidence in subsequent trials (Tsunada et al., 2019; Urai et al., 2019).

Theoretical and experimental evidence suggests that the patterns of stimulus dependence of choice-related signals may be highly informative about the mechanisms of perceptual decision-making. Activity-choice covariations have been characterized in terms of the structure of noise correlations and of feedforward and feedback weights (Shadlen et al., 1996; Cohen and Newsome, 2009; Nienborg and Cumming, 2010; Haefner et al., 2013; Cumming and Nienborg, 2016). Stimulus dependencies may be inherited from the dependence of noise correlations on the stimulus (Kohn and Smith, 2005; Ponce-Alvarez et al., 2013), or from decision-related feedback signals (Bondy et al., 2018). Experimental (Nienborg and Cumming, 2009; Cohen and Maunsell, 2009; Bondy et al., 2018), and theoretical (Maunsell and Treue, 2006; Wimmer et al., 2015; Haefner et al., 2016; Ecker et al., 2016) work indicates that top-down modulations of sensory responses play an important role in the decision-making process. In particular, feedback signals are expected to show cell-specific stimulus dependencies associated with the tuning properties (Lange and Haefner, 2017). Different coding theories attribute different roles to the feedback signals, e. g., conveying predictive errors (Rao and Ballard, 1999) or prior information for probabilistic inference (Lee and Mumford, 2003; Fiser et al., 2010; Haefner et al., 2016; Tajima et al., 2016). Accordingly, characterizing the patterns of stimulus dependence in activity-choice covariations in association with the tuning properties of the cells is expected to provide insights into the particular role of feedback signals and may help to discriminate between alternative proposals. Such an analysis would require simultaneous recordings of populations of neurons tiling the space of receptive fields, and the joint characterization of noise correlations and tuning properties. Although this was beyond the scope of this work, we have shown that our refined methods have a sufficient statistical power to identify a nontrivial structure of stimulus-dependent choice-related signals. The characterization of how the patterns of stimulus dependence vary across brain areas, across cells with different tuning properties, or for different types of sensory stimuli promises to provide further insights about the mechanisms of perceptual decision-making.

While we here analyzed single cell recordings, our conclusions hold for any type of recordings used to study activity-choice covariations. This spans the range from single units (Britten et al., 1996), multiunit activity (Sanayei et al., 2018), and measurements resulting from different imaging techniques at different spatial scales like intrinsic imaging or fMRI (Choe et al., 2014; Thielscher and Pessoa, 2007; Runyan et al., 2017; Michelson et al., 2017). Given the increasing availability of population recordings, larger number of trials due to chronic recordings, and the advent of stimulation techniques to help to discriminate the origin of the choice-related signals (Cicmil et al., 2015; Tsunada et al., 2016; Yang et al., 2016; Lakshminarasimhan et al., 2018; Fetsch et al., 2018; Yu and Gu, 2018), we expect our tools to help gain new insights into the mechanisms of perceptual decision-making.

## 4 Methods

We here describe the derivations of the CP analytical solutions, our new methods to analyze stimulus dependencies in choice-related responses, and describe the data set from Britten et al. (1996) in which we test the existence of stimulus dependencies.

### 4.1 An exact CP solution for the threshold model

We first derive our analytical CP expression valid in the presence of informative stimuli, decision-related feedback, and top-down sources of activity-choice covariation, such as prior bias, trial-to-trial memory, or internal state fluctuations. We follow Haefner et al. (2013) and assume a threshold model of decision making, in which the choice *D* is triggered by comparing a decision variable *d* with a threshold *θ*, so that if *d* > *θ* choice *D* = 1 is made, and *D* = −1 otherwise. The identification of the binary choices as *D* = ±1 is arbitrary and an analogous expression would hold with another mapping of the categorical variable. The choice probability (Britten et al., 1996) of cell *i* is defined as

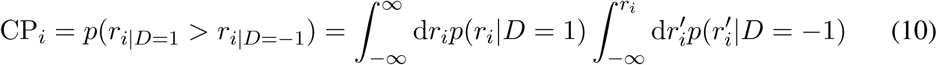

and measures the separation between the two choice-specific response distributions *p*(*r*_*i*_|*D* = −1) and *p*(*r*_*i*_|*D* = 1). It quantifies the probability of responses to choice *D* = 1 to be higher than responses to *D* = −1. If there is no dependence between the choice and the responses this probability is CP = 0.5. To obtain an exact solution of the CP we assume that the distribution *p*(*r*_*i*_, *d*) of the responses *r*_*i*_ of cell *i* and the decision variable *d* can be well approximated by a bivariate Gaussian. Under this assumption, following Haefner et al. (2013) (see their Supplementary Material) the probability of the responses for choice *D* = 1 follows the distribution

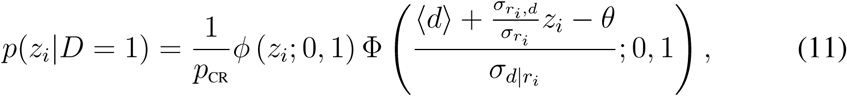

where a more parsimonious expression is obtained using the z-score 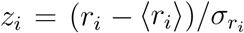. This distribution is a skew-normal (Azzalini, 1985), where *ϕ*(·; 0, 1) is the standard normal distribution with zero mean and unit variance, and Φ(·; 0, 1) is its cumulative function. Furthermore, 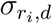 is the covariance of *r*_*i*_ and 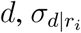 is the conditional standard deviation of *d* given *r*_*i*_, and the probability of *D* = 1 is

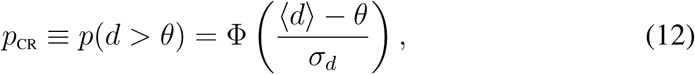

which determines the ratio of the choices or the rate of each choice over trials (the choice ratio or choice rate, respectively). The choice *D* = −1 could equally be taken as the choice of reference, resulting in an analogous formulation. Intuitively, *p*_CR_ increases when the mean of the decision variable ⟨*d*⟩ is higher than the threshold *θ*, and decreases when its standard deviation *σ*_*d*_ increases. Consistently, for an uninformative stimulus, *p*_CR_ = 0.5. Eq. 11 can be synthesized in terms of *p*_CR_ and the correlation coefficient 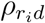, which was named by Pitkow et al. (2015) *choice correlation* (CC_*i*_). In particular, defining 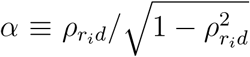 and 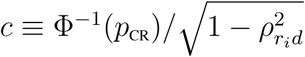

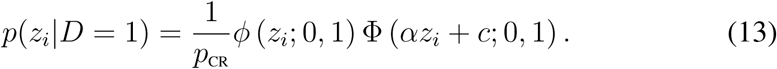

The CP is completely determined by *p*(*z*_*i*_|*D* = −1) and *p*(*z*_*i*_|*D* = 1), and these distributions depend only on *p*_CR_ and the correlation coefficient 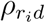. Plugging the distribution of Eq. 13 into the definition of the CP (Eq. 10) an analytical solution is obtained:

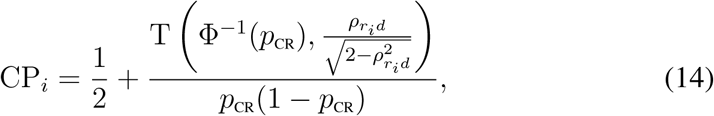

where T is the Owen’s T function (Owen, 1956). In section S1 of the Supplementary Material we provide further details of how this expression is derived. For an uninformative stimulus (*p*_CR_ = 0.5), the function T reduces to the arctangent and the exact result obtained in Haefner et al. (2013) is recovered. The dependence on 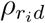 can be intuitively understood because under the Gaussian assumption the linear correlation captures all the dependence between the responses and the decision variable *d*. The dependence on *p*_CR_ reflects the influence of the threshold mechanism, which maps the dependence of *r*_*i*_ with *d* into a dependence with choice *D* by partitioning the space of *d* in two regions.

While Eq. 14 provides an exact solution of the CP, in the Results section we present and mostly focus on a linear approximation to understand how the stimulus content modulates the choice probability. This approximation is derived (see S1 in Suppl. Material) in the limit of a small 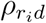, which leads to CPs close to 0.5 as usually measured in sensory areas (Nienborg et al., 2012). However, as we show in the Results and further justify in the Suppl. Material this approximation is robust for a wide range of 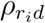 values. The linear approximation relates the choice probability to the Choice Triggered Average (CTA) (Haefner, 2015; Chicharro et al., 2017), defined as the difference of the mean responses for each choice. The mean response of cell *i* in trials where *D* = 1 is made is

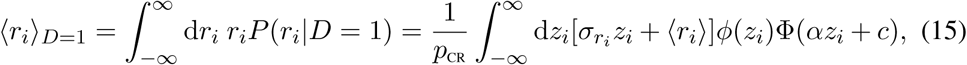

where the last equality holds for *P* (*r*_*i*_|*D* = 1) having the form of Eq. 13. This formula can be analytically solved to obtain a closed-form expression for the CTA

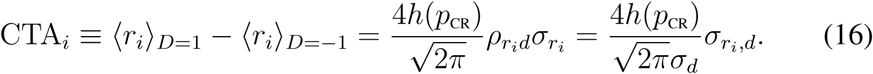

Furthermore, the fact that *D* is a binary variable, without any other assumption about the distribution of the responses, implies that

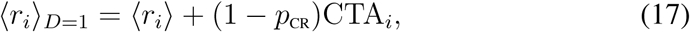

and similarly for ⟨*r*_*i*_⟩ _*D*=−1_ substituting 1 − *p*_CR_ by −*p*_CR_. That is, it is the conditional mean of the less likely choice the one that departs more from ⟨*r*_*i*_⟩. The detailed form of the linear approximation of the CP in terms of the CTA is described in the Results section.

### 4.2 Neuronal data

To study stimulus dependencies in the relationship between the responses of sensory neurons and the behavioral choice we analyzed the data from Britten et al. (1996) publicly available in the Neural Signal Archive (www.neuralsignal.org). In particular, we analyzed data from file nsa2004.1, which contains single unit responses of macaque MT cells during a random dot discrimination task. This file contains 213 cells from three monkeys. We also used file nsa2004.2, which contains paired single units recordings from 38 sites from one monkey. For the single unit recordings, the direction tuning curve of each neuron was used to assign a preferred-null axis of stimulus motion, such that opposite directions along the axis yield a maximal difference in responsiveness (Bair et al., 2001). For paired recordings, the direction of stimulus motion was selected based on the direction tuning curve of the two neurons and the criterion used to assign it varied depending on the similarity between the tuning curves. For cells with similar tuning, a compromise between the preferred directions of the two neurons was made. For cells with different tuning, the axis were chosen to match the preference of the most responsive cell. To minimize the influence in our analysis of the direction of motion selection, we only analyzed the most responsive cell from each site. Accordingly, our initial data set consisted in a total of 251 cells. The same qualitative results were obtained when limiting the analysis to data from nsa2004.1 alone. Further criteria regarding the number of trials per each stimulus level were used to select the cells. As discussed below, if not indicated otherwise, we present the results from 107 cells that fulfilled all the criteria required.

### 4.3 Analysis of stimulus-dependent choice probabilities

Our analysis of choice probabilities stimulus dependencies is based on examining the patterns in the CP(*p*_CR_) profile as a function of the probability *p*_CR_ ≡ *p*(*D* = 1). We here describe how these profiles are constructed, the surrogates-based method used to assess the significance of stimulus dependencies, and the clustering analysis used to identify different stimulus dependence patterns. Matlab functions to calculate weighted average CPs, to obtain CP profiles, and to generate surrogates consistent with the null hypothesis of a constant CP are to be available at http://www2.bcs.rochester.edu/sites/haefnerlab.

#### 4.3.1 Profiles of CP as a function of the ratio of choices

We constructed CP(*p*_CR_) profiles instead of CP(*s*) profiles based on the prediction from the theoretical threshold model of the modulatory factor *h*(*p*_CR_). We estimated the *p*_CR_ value associated with each random dots coherence level using the psychophysical function for each monkey separately. For each coherence level, we calculated a CP value if at least 15 trials were available in total, and at least 4 for each choice. In the original analysis of Britten et al. (1996) stimulus dependencies CP(*s*) were examined averaging across cells the CP at each coherence level. This analysis did not separate the within-cell stimulus dependencies CP(*s*) from variability due to changes in choice probabilities across cells. In particular, in the data set the stimulus levels presented vary across cells, which means that for each coherence level the average CP does not only reflect any potential stimulus dependence of the CP but also which subset of cells contribute to the average at that level. Therefore, we binned the range of *p*_CR_ in a way that for each cell at least one stimulus level mapped to each bin of *p*_CR_. We here present the results using five bins defined as [0 − 0.3, 0.3 − (0.5 −*ε*), (0.5 −*ε*) − (0.5+*ε*), (0.5+*ε*) −0.7, 0.7 − 1], where *ε* was selected such that only trials with the uninformative (zero coherence) stimulus were comprised in the central bin. Results are robust to the exact definition of the bins. We selected larger bins for highly informative stimulus levels for two reasons. First, the stimulus levels used in the experimental design do not uniformly cover the range of *p*_CR_, there are more stimulus levels corresponding to *p*_CR_ values close to *p*_CR_ = 0.5. Second, the CP estimates are worse for highly informative stimuli. In particular, the standard error of the CP estimates depends on the magnitude of the CP itself (Bamber, 1975; Hanley and McNeil, 1982) but for small |CP − 0.5| can be approximated as

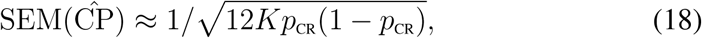

where *K* is the number of trials. The product *p*_CR_(1 − *p*_CR_) is maximal at *p*_CR_ = 0.5 and decreases quadratically when *p*_CR_ approximates 0 or 1. Furthermore, in the data set the number of trials *K* is higher for stimuli with low information, while most frequently *K* = 30 for highly informative stimuli. We used these estimates of the 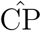 error to combine the CPs of *M*_*k*_ different stimulus levels assigned to the same bin *k* of *p*_CR_. The average CP(*p*_CR,*k*_) for bin *k* was calculated as 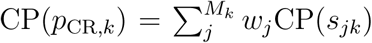 with normalized weights proportional to 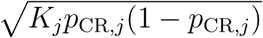. A full profile CP(*p*_CR_) could be constructed for 107 cells, while for the rest a CP value could not be calculated for at least one of the bins because of the criteria on the number of trials. Together with the profile CP(*p*_CR_) we also obtained an estimate of its error as a weighted average of the errors, which corresponds to

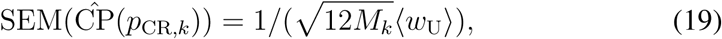

where ⟨*w*_U_⟩ is the average of the unormalized weight 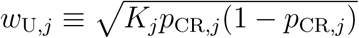. Following this procedure we can iteratively calculate weighted averages of the CPs across different sets. In particular, we used this same type of average to obtain averaged CP(*p*_CR_) profiles across cells. Importantly, in contrast to the analysis of Britten et al. (1996), we previously separated the cells into two groups, with a positive or negative average CP−0.5 value, given that the effect of *h*(*p*_CR_) predicts an inverse modulation by *p*_CR_.

#### 4.3.2 Surrogates to test the significance of CP stimulus dependencies

Given a certain average profile CP(*p*_CR_), we want to assess whether the pattern observed is compatible with the null hypothesis of a constant CP value for all *p*_CR_ values. In particular, because the error of the CP estimates is sensitive to the number of trials *K* and to *p*_CR_ (Eq. 18), we want to discard that any structure observed is only a consequence of changes of *K* and *p*_CR_ across the bins used to calculate the CP(*p*_CR_) profiles. For this purpose, we developed a procedure to build surrogate data sets compatible with the hypothesis of a flat CP(*p*_CR_) and that preserves at each stimulus level the number of trials for each choice. The surrogates are built shuffling the trials across stimulus levels to destroy any stimulus dependence of the CP. However, because the responsiveness of the cell changes across levels according to its direction tuning curve, responses need to be normalized before the shuffling. Kang and Maunsell (2012) showed that, to avoid underestimating the CPs, this normalization should take into account that mean responses at each level are determined by the conditional mean response for each choice and also by the ratio of choices. Under the assumption of a constant CP, they proposed an alternative z-scoring, which estimates the mean and standard deviation correcting for the different contribution of trials corresponding to the two choices (see section S2 in the Suppl. Materials for details of their method).

We applied the z-scoring of Kang and Maunsell (2012) to pool the responses within an interval of stimulus levels with low information, preserving only the separation of trials corresponding to each choice. We selected the interval from −1.6% to 1.6% of coherence values, which comprises a third of the informative coherence levels used in the experiments. Because these stimuli have low information they lead to *p*_CR_ values close to *p*_CR_ = 0.5 and hence we can approximate the CP as constant within this interval. The fact that the factor *h*(*p*_CR_) is almost constant around *p*_CR_ = 0.5 (see Figure 2A) further supports this approach. We used this pool of neural responses to sample responses for all stimulus levels in the surrogate data set. For each stimulus level of the surrogate data, the number of trials for each choice was preserved as in the original data. In these surrogates, apart from random fluctuations, any structure in the CP(*p*_CR_) profiles can only be produced by the changes in *K* and *p*_CR_ across bins. To test the existence of significant stimulus dependencies in the original CP(*p*_CR_) profiles we calculated the differences ΔCP_*k*_ = CP(*p*_CR,*k*+1_) − CP(*p*_CR,*k*_) for the bins *k* = 1, …, 4. To test for an asymmetric pattern with respect to *p*_CR_ = 0.5 the average of ΔCP_*k*_ across bins was calculated. To test for a symmetric pattern the sign of the difference was flipped for the bins corresponding to *p*_CR_ < 0.5 before averaging. When testing for a pattern consistent with the modulation predicted by the threshold model, the shape was inverted for cells with average CP lower than 0.5. The same procedure was applied to each surrogate CP(*p*_CR_) profile. We generated 8000 surrogates and estimate the p-value as the number of surrogates for which the average ΔCP was higher than for the original data.

#### 4.3.3 Clustering analysis

We used nonparametric and parametric clustering analysis to examine the patterns in the CP(*p*_CR_) profiles beyond the stereotyped shape *h*(*p*_CR_) predicted from the threshold model. We first used nonparametric *k*-mean clustering for an exploratory analysis of which patterns are more common among the 107 cells for which a complete CP(*p*_CR_) profile could be constructed. The clustering was implemented calculating cosine distances between vectors defined as CP(*p*_CR_)−0.5. The selection of this distance is consistent with the prediction of the threshold model that a different pattern is expected for cells with a CP higher or lower than 0.5. We examined the patterns associated with the clusters as a function of the number of clusters to identify robust patterns of dependence (see Figure S1B,C). We then focused on a symmetric and an asymmetric pattern of CP(*p*_CR_) with respect to *p*_CR_ = 0.5. To better interpret these two clusters we complemented the analysis with a parametric clustering approach in which a symmetric and asymmetric template were a priori selected to cluster the CP(*p*_CR_) profiles. To assess the significance of the CP(*p*_CR_) patterns we repeated the same clustering procedure for surrogate data generated as described above. See section S3 in the Supplementary Material for a more detailed description of the construction, visualization, and significance assessment of the CP(*p*_CR_) patterns.

### 4.4 The effect of response gain fluctuations on choice probabilities

Consider the classic feedforward encoding/decoding model (Fig. 1B, Shadlen et al., 1996; Haefner et al., 2013) in which a population of sensory responses, **r** = (*r*_1_, …, *r*_*n*_), is read out into a decision variable

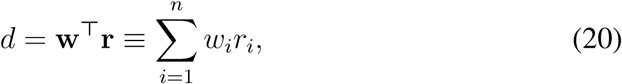

where **w** are the read-out weights. The categorical choice *D* is made by comparing *d* to a threshold *θ*. We model the responses as *r*_*i*_ = *f*_*i*_(*s*)+*ξ*_*i*_, with tuning functions **f** (*s*) = (*f*_1_(*s*), …, *f*_*n*_(*s*)) and a covariance structure Σ of the neuron’s intrinsic variability *ξ*_*i*_. The read-out weights **w** selected such that *d* represents an optimal estimate ŝ of the sensory stimulus *s* are determined by the noise covariance matrix and the tuning curves (Haefner et al., 2013; Pitkow et al., 2015; Moreno-Bote et al., 2014). In particular, the decoder is optimal if the weights are optimized to the structure of the covariance matrix at the decision boundary (uninformative) stimulus *s*_0_:

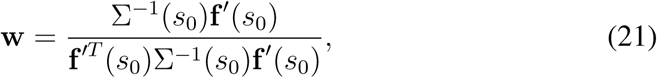

where **f**′(*s*_0_) and Σ(*s*_0_) are the derivative of the tuning curves and the noise covariance matrix, respectively, for *s* = *s*_0_.

While these read-out weights are optimized according to the noise structure at the decision boundary, the variability and co-variability of neural responses are stimulus dependent (Lee et al., 1998; Josić et al., 2009). Neural responses with Poisson-like statistics have a variance proportional to the mean firing rate. Furthermore, excitability gain fluctuations lead to a squared dependence of the variance on the firing rate (Pillow and Scott, 2012; Goris et al., 2014; Ecker et al., 2016). Goris et al. (2014) showed that in MT gain fluctuations can account for more than 75% of the variance in the responses to sinusoidal gratings. These excitability fluctuations appear if the tuning curve is modulated by trial-to-trial fluctuations such that, for cell *i* in trial *k, f*_*ik*_(*s*) = *g*_*k*_*f*_*i*_(*s*), where *g*_*k*_ is a gain modulatory factor. The covariance matrix can be partitioned as

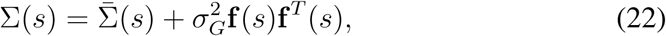

where 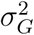 is the variance of the gain *g* and 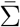 is the covariance not associated with the gain, which comprises the Poisson-like variability. For simplicity, we only considered the case in which a global gain fluctuation affects the response of the whole population. More generally, the magnitude of the gain may vary across cells, as well as the degree to which the gain co-fluctuates across cells. Furthermore, we also approximated 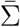 as stimulus independent to focus on how gain fluctuations produce choice probabilities stimulus dependencies. For this purpose, we studied how the CP is affected when the optimal read-out weights of Eq. 21 are used to construct the decision variable but the responses have a covariance structure Σ(*s*) different from Σ(*s*_0_), rendering the weights suboptimal for *s* ≠ *s*_0_. We used a linear approximation on *s* – *s*_0_ of the stimulus-dependent covariance matrix in Eq. 22 and determined how these stimulus dependencies affect the CP (see section S4 in the Suppl. Material for details). In the Results section we study the relation between the strength of the gain fluctuations 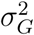, the magnitude of the CP, and the CP dependence on *p*_CR_.

### 4.5 Generalized Linear Models modeling the interaction between stimulus and choice predictors

The existence of a stimulus dependence of the CP indicates that the activity-choice covariation is modulated by the stimulus level. When modeling the neural responses with Generalized Linear Models (GLMs), it is common to treat the sensory stimulus and the choice as separate predictors with no interaction, using a single parameter (or kernel) to account for the effect of the choice (Park et al., 2014). We examined the improvement in the fit of the neural responses when using a GLM model in which several parameters model the influence of the choice, associated with different levels of the stimulus. In more detail, we fitted nested Poisson GLM models to the spike counts of each trial, progressively increasing the number of parameters. In the simplest model, a constant rate is assumed. We then considered models with the stimulus level as a predictor. In particular, a fourth order polynomial of the coherence level is used to model the firing rate. We subsequently added the choice as a separate single predictor. Finally, we considered models with multiple parameters (*N*_*c*_ = 2, 3) associated with the choice predictor, each accounting for the influence of the choice in a certain range of *p*_CR_. The full model has the form Eq. 9. For the models with multiple choice parameters, the subsets of *p*_CR_ associated with each choice-parameter level were determined based on the CP(*p*_CR_) profile. In particular, for *N*_*c*_ > 1, the existence of nonmonotonic CP(*p*_CR_) profiles, such as the symmetric pattern around *p*_CR_ = 0.5, indicated that it would be suboptimal to tile the domain of *p*_CR_ with *N*_*p*_ bins and assign a different choice-parameter level to each bin. Accordingly, we first estimated the CP(*p*_CR_) profile of each cell and then used *k*-mean clustering with an Euclidean distance to cluster the components of CP(*p*_CR_), corresponding to the bins of *p*_CR_, into *N*_*c*_ subsets. A different GLM choice-parameter *b*_*j*_ was then assigned to each choice-parameter level *j* = 1, …*N*_*c*_.

We compared the predictive power of the nested models using cross-validation. To avoid that the choice-parameters fitted were affected by the ratio of trials with each choice, we matched the number of trials of each choice used to fit the model at each choice-parameter level. In more detail, we first merged in two pools, one for each choice separately, the trials of all stimulus levels assigned to the same choice-parameter level. We then determined the number of trials from each pool to be included in the fitting set as an 80% of the trials available in the smallest pool, hence matching the number of trials selected from each choice. The remaining trials were left for the testing set. This procedure was repeated for each choice-parameter level and a GLM model was fitted on the fitting set obtained combining the selected trials for all levels. This random separation between fitting and testing data sets was repeated 50 times and the average predictive power was calculated. Performance was then quantified comparing the increase in the likelihood of the data in the testing set with respect to the likelihood of the null model which assumes a constant firing rate (*L*_0_). To determine if incorporating the choice as a predictor improved the prediction, we examined the likelihood relative increase (LRI) defined as the ratio of the likelihood increase *L*(*choice, stimulus*) − *L*(*stimulus*) and the increase *L*(*stimulus*) − *L*_0_. For the models allowing for multiple choice parameters we selected the most predictive model from *N*_*c*_ = 2, 3. To evaluate the improvement when considering stimulus-dependent choice influences we compared the **LRI** obtained for the models with a single or multiple parameters.

## Acknowledgments

This work was supported by the BRAIN Initiative (Grants No. R01 NS108410 and No. U19 NS107464 to S.P.), the Fondation Bertarelli, and the CRCNS program under R01 EY028811 to R.M.H. We thank K.H. Britten, W.T. Newsome, M.N. Shadlen, S. Celebrini, and J.A. Movshon for making their data available in the Neural Signal Archive (http://www.neuralsignal.org/), and W. Bair for maintaining this database.

## Conflict of Interest

The authors declare no competing financial interests.

## Authors Contributions

DC: Conceptualization, Data curation, Formal analysis, Investigation, Methodology, Visualization, Writing – original draft. SP: Funding acquisition, Supervision, Writing – review and editing. RMH: Conceptualization, Formal analysis, Funding acquisition, Methodology, Supervision, Writing – review and editing.

## Supplementary Material

### S1 The analytical expression of choice probability

We here provide details of how the CP analytical expression of Eq. 14 is obtained from the definition of Choice Probability (Eq. 10) when the probability of the responses for each choice has the form of Eq. 13, derived from the threshold model. Plugging the distribution of Eq. 13 into the definition of the CP we get

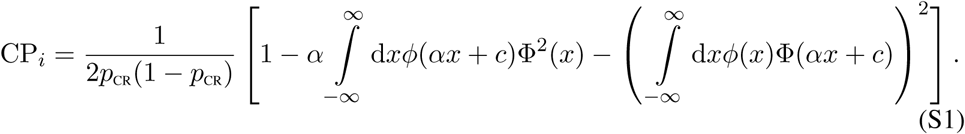

This expression is derived analogously to Eq. S1.2 in Haefner et al. (2013), and generalizes the case examined there, which corresponds to *c* = 0. To solve this expression we use some results involving integrals of normal distributions:

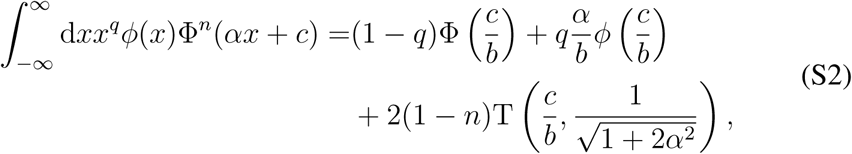

where 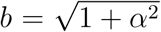 and T is the Owen’s T function (Owen, 1956). The equality above is valid for the cases *q* = 0, *n* = 1, 2, and *q* = 1, *n* = 1, which we used to derive the expressions of the CP and CTA. Using the equality for *q* = 0, *n* = 1, 2 into Eq. S1 we obtain the CP expression of Eq. 14. On the other hand, the case with *q* = 1, *n* = 1 allows deriving the expression of the CTA (Eq. 16) from Eq. 15.

The CP linear approximation of Eq. 2 is generically valid when the activity-choice covariations are well captured by the linear dependence between the responses and the choice. We here only present a restricted derivation, specifically from the exact CP solution resulting from the threshold model. It can be checked that the same approximation follows for example from the exact solution of the CP obtained when taking the conditional distributions *p*(*r*_*i*_|*D*) to be Gaussians (Dayan and Abbot, 2001; Carnevale et al., 2013) and not skew normals (Eq. 13) like for the threshold model. Expanding Eq. 14 in terms of 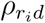 we get a polynomial approximation

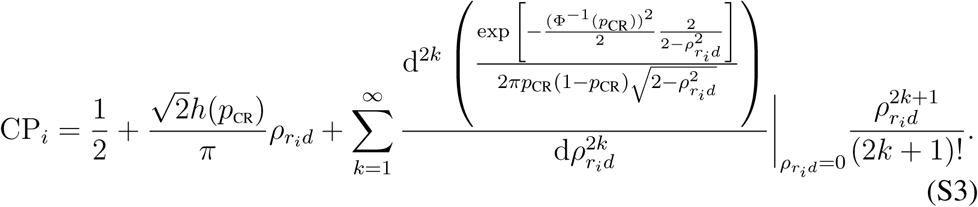

This expansion contains only odd order terms, producing a symmetry of CP−0.5 with respect to the sign of 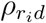. This explains why Haefner et al. (2013) found that the linear approximation was accurate for a wide range of 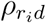 values, since the choice correlation needs to be high so that the contribution of 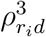 is relevant. Up to order 3

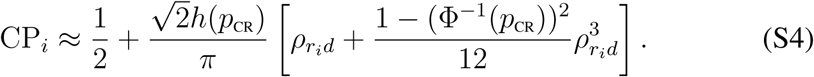

Since Φ^−1^(0.5) = 0, for *p*_CR_ values for which |1 − (Φ^−1^(*p*_CR_))^2^| < 1 the third order term makes a smaller contribution than for the uninformative case. This is true for 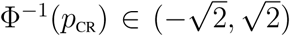, which leads to *p*_CR_ ∈ (0.08, 0.92). This means that the linear approximation is expected to be an even better approximation in this range than for *p*_CR_ = 0.5. Furthermore, for (Φ^−1^(*p*_CR_))^2^ < 1 the third order contribution is positive, so that for the *p*_CR_ values fulfilling this constraint, *p*_CR_ ∈ (0.16, 0.84), the linear approximation is expected to underestimate the CP. The range of *p*_CR_ in which the linear approximation underestimates or overestimates the CP can be seen in Figure 2 of the main article.

### S2 The relation between the weighted average CP and the grand CP of z-scored responses

We here review how z-scoring is used to calculate a grand CP (Britten et al., 1996) pooling the responses across stimulus levels. Kang and Maunsell (2012) provided a qualitative explanation of why the standard z-scoring of the responses leads to an underestimation of the CP (see their Figure 1) and proposed a corrected z-scoring to pool the responses across stimulus levels. We use the expression of the CP in terms of the CTA (Eq. 2) to understand the connection between these procedures and the calculation of a weighted average of the CP values across stimulus levels. We show that the underestimation of the CP with standard z-scoring is associated with an improper use of unormalized weights, while the corrected z-scoring corresponds to a particular choice of normalized weights. We will use *z*_*s*_ to refer to the standard z-scored responses and *z*_*c*_ to refer to responses with the corrected z-scoring. In more detail, for each stimulus level *s*, the standard z-scoring corresponds to *z*_*s*_ = (*r* − *µ*_*r*|*s*_)/*σ*_*r*|*s*_. The corrected z-scoring of Kang and Maunsell (2012) balances the weight of the choice conditional means and variances in the normalization and considers 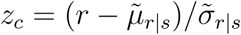 with

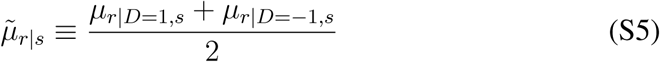

and

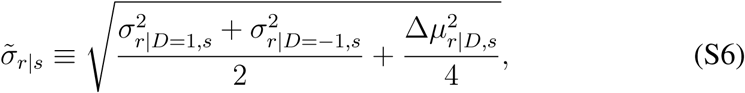

where Δ*µ*_*r*|*D,s*_ is the CTA for the stimulus level *s*. As discussed in the Methods, *µ*_*r*|*D*=1,*s*_ = *µ*_*r*|*s*_ +(1−*p*CR)Δ*µ*_*r*|*D,s*_ (Eq. 17) and *µ*_*r*|*D*=−1,*s*_ = *µ*_*r*|*s*_ −*p*CRΔ*µ*_*r*|*D,s*_. Accordingly, 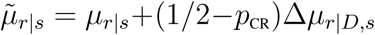. On the other hand, 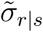 only introduces a second-order correction with respect to the standard normalization with *σ*_*r*|*s*_. In particular, given the skew-normal distributions (Eq. 13) resulting from the threshold model, both 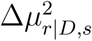 and 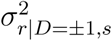 depend quadratically in the strength of the activity-choice covariations, as determined by the choice correlation (Arnold and Beaver, 2000; Azzalini, 2005). Accordingly, to understand the difference between the standard and corrected z-scoring, we neglect the correction in 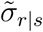 and focus on how demeaning with 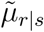 differs from using *µ*_*r*|*s*_. Equivalently to Eq. 2, the grand CP from the pooled z-scored responses can be approximated as

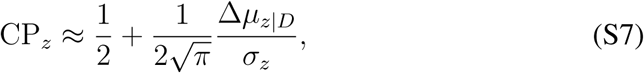

where we drop the cell index *i* for simplicity. In relation to the main article, we here use a different notation such that 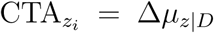 and var *z*_*i*_ = *σ*_*z*_. Furthermore, in this section we explicitly indicate which measures are calculated for a fixed stimulus, while a CP or Δ*µ*_*z*|*D*_ without a stimulus subindex refers to the grand measures. In the case of *p*_CR_, it is defined as *p*_CR_ ≡ *p*(*D* = 1|*s*) as before, although previously the dependence on the fixed stimulus was implicit. In relation to the grand CP, the CTA is calculated from the distributions *p*(*z*|*D* = 1), *p*(*z*|*D* = −1), and *σ*_*z*_ from *p*(*z*). When the stimulus level is fixed, the same linear approximation holds for CP(*s*) in terms of Δ*µ*_*z*|*D,s*_ and *σ*_*z*|*s*_, calculated from *p*(*z*|*D* = 1, *s*), *p*(*z*|*D* = −1, *s*), and *p*(*z*|*s*), respectively. Given that at all stimulus levels *σ*_*z*|*s*_ = 1, also *σ*_*z*_ = 1, and hence the CP can be understood as reflecting the CTA of the normalized responses.

We now use the decomposition of the mean as the average of the conditional means across stimulus levels to express Δ*µ*_*z*|*D*_ in terms of Δ*µ*_*z*|*D,s*_. Given *µ*_*r*|*D*=1,*s*_ = *µ*_*r*|*s*_+(1−*p*_CR_)Δ*µ*_*r*|*D,s*_, we obtain 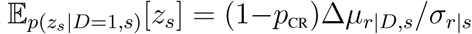. Given 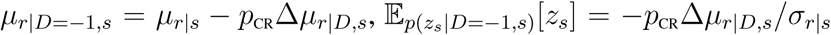. Accordingly, with the standard z-score

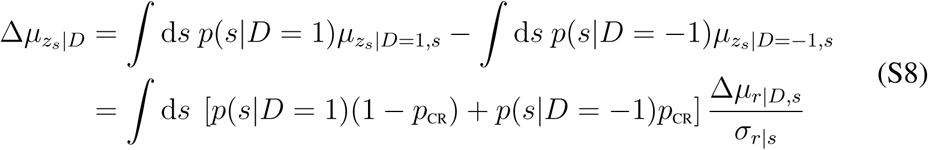

Taking into account that Δ*µ*_*r*|*D,s*_/*σ*_*r*|*s*_ corresponds in the linear approximation to 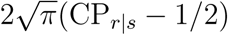, Eq. S8 can be expressed for the CP as

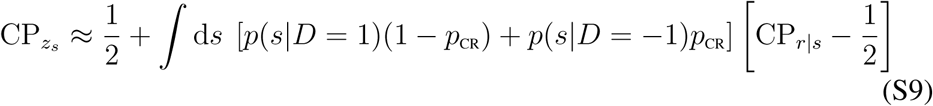

We can now understand why the standard z-scoring is inadequate to estimate the grand CP. In particular, 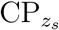 corresponds to a weighted average of the CP values for each stimulus level, with weights defined as 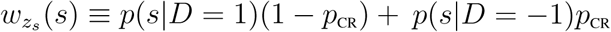. However, these weights are not normalized, because 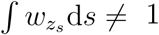.

Oppositely, the grand CP calculated with the corrected z-scoring corresponds to a weighted average with normalized weights. Following the same procedure to decompose the mean as an average of conditional means, given 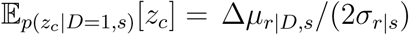 and 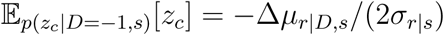 we obtain

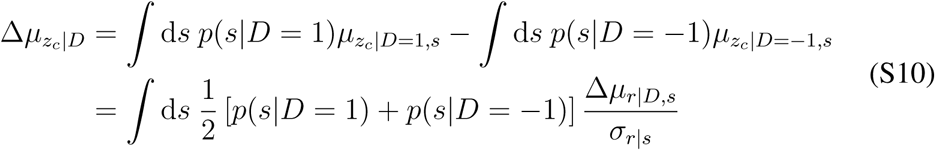

and

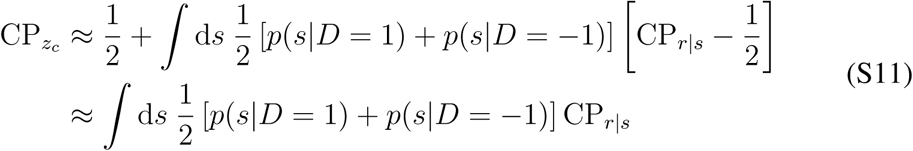

where the weighted average has weights 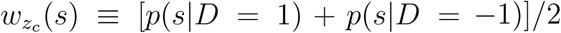, which are properly normalized to 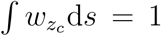. If across all stimulus levels the rate of the choices is balanced, i. e. if *p*(*D* = 1) = *p*(*D* = −1), then the weights simplify to 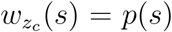, that is, the CPs are averaged according to the relative number of trials available for each stimulus level given the experimental settings and recording contingencies.

Kang and Maunsell (2012) already suggested to use an explicit weighted average of the CP values as an alternative to the grand average calculated with the corrected z-scores. The analysis above shows that in fact the use of z-scores corresponds in the linear approximation to a specific weighted average, with weights 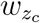. These weights can be reexpressed as

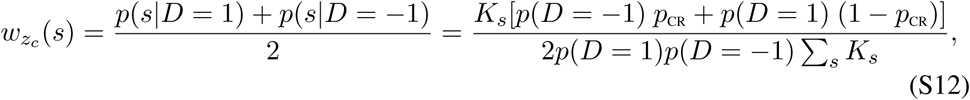

where ∑ _*s*_ *K*_*s*_ is the total number of trials and *K*_*s*_ the trials with stimulus *s*. Because the denominator is common to all stimulus levels, the weights are proportional to *K*_*s*_[*p*(*D* = −1) *p*_CR_ + *p*(*D* = 1) (1 − *p*_CR_)], in contrast to the weights inversely proportional to the standard error of the CP estimates (Eq. 18), which are proportional to 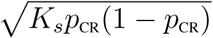. This suggests that, although the corrected z-scoring does not induce any systematic bias, the weighted average adapted to the estimation errors may be preferable. For this reason, we used this latter type of weighted average when combining CP values across stimulus levels or across cells, and only use the pooling procedure to construct the surrogates for significance testing. In that case, it is the CP variability across stimulus levels what is evaluated, and the number of trials generated for each choice at each stimulus level is preserved to avoid that the original and surrogate CPs differ due to estimation issues.

### S3 Clustering analysis

We here provide further details about the alternative procedures used to cluster the CP(*p*_CR_) profiles, about the visualization of the clusters, and about how to assess the significance of the CP(*p*_CR_) patterns. As a first step, we implemented a non-parametric *k*-mean clustering analysis to cluster the CP(*p*_CR_) profiles of the 107 cells for which a full profile could be constructed. We started using *C* = 2 clusters (Fig. 4A) and found that this nonparametric approach, when using the cosine distance, recovered qualitatively the same patterns obtained when separating a priori the cells into cells with an average CP higher or lower than 0.5 (Fig. 3A). From the patterns of the two clusters only the one of cells with an average CP higher than 0.5 was found significant (see below for details on the significance analysis). Given this difference in significance, we subsequently increased the number of clusters in two alternative ways. In a first approach, we a priori separated the cells with an average CP higher or lower than 0.5 and continued the clustering analysis separately for these two groups. Fig. S1A shows the obtained subclusters with *C* = 2 for the two groups separately. As a second approach, we increased the number of clusters to *C* = 3 without any previous separation in two groups. The resulting clusters (Fig. S1B) indicated that the separation of the two subclusters for the cells with average CP higher than 0.5 naturally appears without enforcing the separation. Increasing the number of clusters without any a priori separation provided evidence that the two main patterns for cells with average CP higher than 0.5 are robust and still contain a substantial portion of the cells even for *C* = 6 (Fig. S1C). We therefore focused our posterior analysis in characterizing the features of these symmetric and asymmetric patterns.

To evaluate the significance of the CP(*p*_CR_) patterns found with the clustering analysis we repeated the same clustering procedure for the surrogate data generated as described in Methods. For each surrogate, each of the *C* clusters found was associated with the most similar original pattern of the ones being tested. For example, in Fig. 4B, when testing the significance of the symmetric and asymmetric patterns for cells with average CP higher than 0.5, each of the two surrogate cluster patterns was assigned to the most similar pattern among the symmetric and asymmetric one. Subsequently, the average of ΔCP_*k*_ across bins was calculated for the original and surrogate profiles as explained in the Methods. The p-value corresponding to each original pattern was calculated from the number of surrogate patterns associated with it for which the average ΔCP_*k*_ was higher.

To visualize the clusters in Figs. 4 and 5 we constructed orthonormal axes using either the vectors corresponding to the center of the clusters or the selected templates, for nonparametric and parametric clustering, respectively. In the case of nonparametric clustering, the x-axis corresponds to the separation between the two initial clusters, and is closely aligned to the departure of the average CP from 0.5. The y-axis was built as a projection orthogonal to the x-axis of the vector connecting the center of the two subclusters. When templates were used, the x-axis corresponds to the template with a constant CP and the y-axis was built as an orthogonal projection of the template with an asymmetric profile (a vector with a positive unit slope).

**Figure S1:**
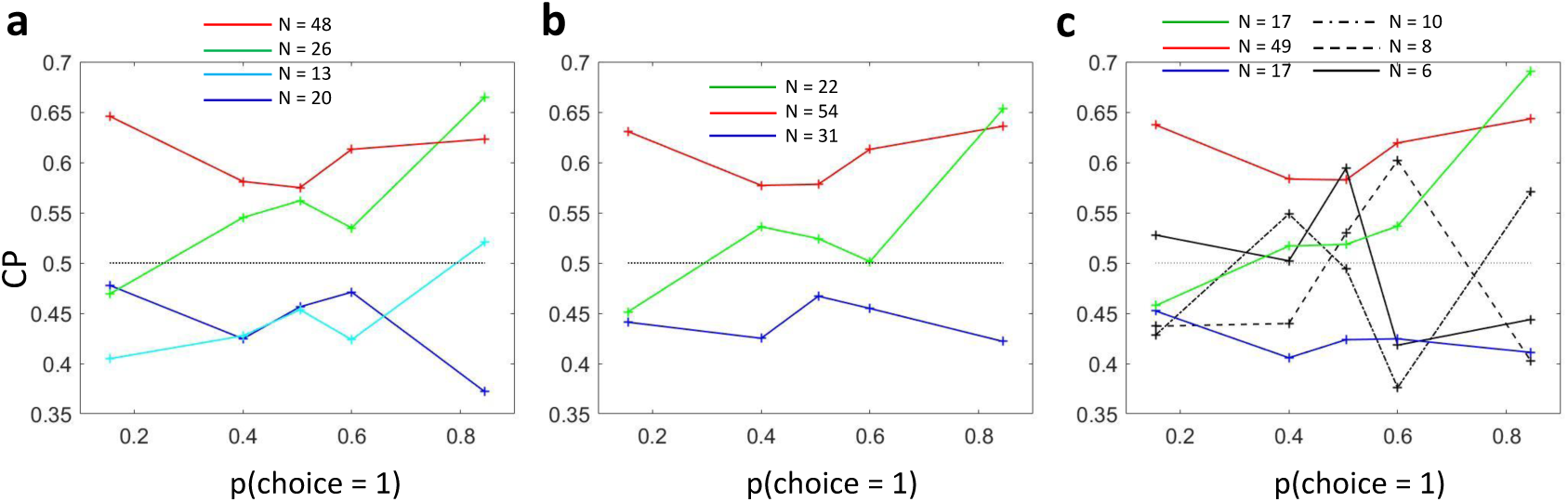
Subclustering of CP(*p*_CR_) dependencies. **a**) Analogous to Fig. 4B but showing also the average profiles for the two subclusters obtained from cells with average CP < 0.5. **b**) Nonparametric *k*-mean clustering with three clusters determined from all cells. **c**) Nonparametric *k*-mean clustering with six clusters determined from all cells. The clusters more similar to the ones of Fig. 4B are correspondingly coloured.

### S4 The effect of gain fluctuations on the CP

We here derive the expression of the CP in terms of the strength of the gain fluctuations 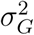, as expressed in Eq. 8 in the main paper. Consider that the decoder used is optimal for the uninformative stimulus at the decision boundary (Eq. 21) and that the covariance matrix is stimulus dependent due to a component induced by gain fluctuations (Eq. 22). Given the linear approximation

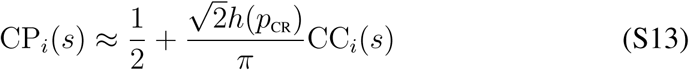

the stimulus dependence of CP_*i*_(*s*) appears through the factor *h*(*p*C_R_) and also through the stimulus dependence of the choice correlation CC_*i*_(*s*). For the threshold model, the choice correlation depends on the read-out weights and on the noise covariance matrix as (Haefner et al., 2013)

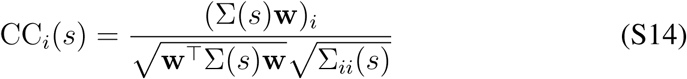

where 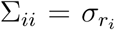 and 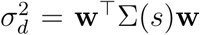. When the stimulus presented is the uninformative *s*_0_, the covariance Σ(*s*_0_) is the one for which the decoder **w** has been optimized. Using the optimal weights (Eq. 21) in Eq. S14 the choice correlation has the form

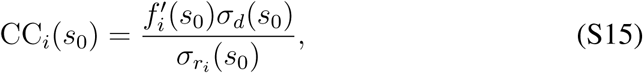

which is proportional to the ratio between the neural threshold of neuron 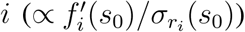 and the behavioral threshold (∝ 1/*σ*_*d*_(*s*_0_)) (Pitkow et al., 2015). The CC_*i*_ depends on the sensitivity of cell *i* itself because the weights are optimized to invert the noise correlations between the responses by Σ−1(*s*_0_). However, if Σ(*s*) is stimulus dependent as in the presence of gain fluctuations (Eq. 22), this cancelation of the effect of the noise covariance only holds for the uninformative stimulus. In a first order approximation, the covariance matrix changes with an stimulus change *s* − *s*_0_ as

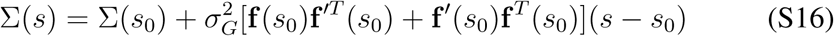

and for this covariance the weights are not completely optimal. We used this expression of Σ(*s*) to derive the form of the terms (Σ(*s*)**w**)_*i*_, **w**^T^Σ(*s*)**w**, and Σ_*ii*_(*s*) which determine the CC_*i*_(*s*) in Eq. S14. Plugging Eq. S16 into Eq. S14, and given the relation between the stimulus content and *p*_CR_ (Eq. 12), the following dependence of the choice correlation on *p*_CR_ is obtained:

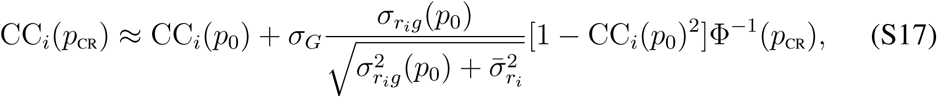

where *p*_0_ corresponds to the uninformative stimulus *s*_0_, and 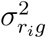 and 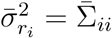 are the part of the variance of cell *i* associated or not with the gain, respectively. That is, 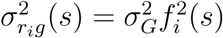 and 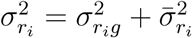.

The expressions of CC_*i*_(*s*_0_) in Eq. S15 and of CC_*i*_(*p*_CR_) in Eq. S17 indicate how the strength of gain fluctuations 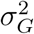 determines the dependence of choice correlation on *p*_CR_. The dependence of CC_*i*_(*s*_0_) on 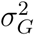 appears through the standard deviation of the cell 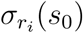. Given the covariance matrix of Eq. 22

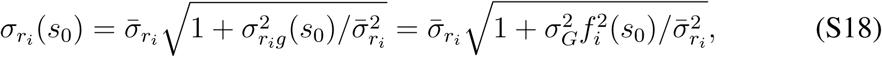

where 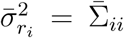. A dependence could also appear through *Σ*_*d*_(*s*_0_) in Eq. S15, but Ecker et al. (2016) have shown that in a model with biologically plausible receptive fields (Maunsell and Treue, 2006) the information about the stimulus (and hence 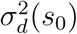) is mostly unaffected by the magnitude of 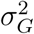 given that the tuning curves and their derivatives vary orthogonally. Accordingly, we considered 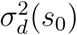 invariant to 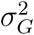 so that the dependence of the choice correlation CC_*i*_(*s*_0_) on 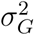 is

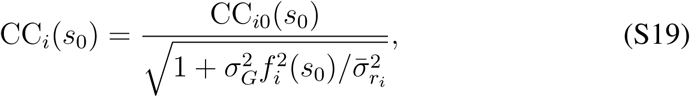

where CC_*i*0_(*s*_0_) is the choice correlation for the case of no gain fluctuations, 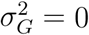. Combining Eqs. S17 and S19 we obtain the expressions of Eq. 8 in the main paper, which show how overall the choice correlation depends on the strength of the gain fluctuations 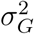 and on *p*_CR_:

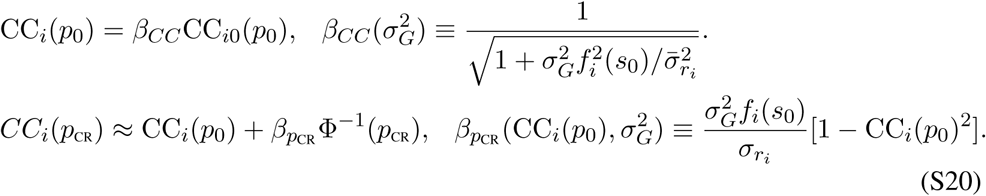

## Footnotes

1 This relation holds for the covariance between any variable *x* and a binary variable *D*, and independently of the convention adopted for the values of *D*: the factor 2 has to be replaced by *a* − *b* in general for *D* = *a, b* instead of *D* = 1, −1.

2 code will be available at Haefner Lab’s webpage

## References

Azzalini, A. (1985). A class of distributions which includes the normal ones. Scandinavian Journal of Statistics, 12:171–178.

Bair, W., Zohary, E., and Newsome, W. T. (2001). Correlated firing in macaque visual area MT: Time scales and relationship to behavior. J. Neurosci., 21(5):1676–1697.

Bamber, D. (1975). The area above the ordinal dominance graph and the area below the receiver operating graph. J. Math. Psychol., 12:387–415.

Bishop, C. M. (2006). Pattern Recognition and Machine Learning. Springer, New York.

Bogacz, R., Brown, E., Moehlis, J., Holmes, P.,, and Cohen, J. D. (2006). The physics of optimal decision making: A formal analysis of models of performance in two-alternative forced-choice tasks. Psychological Review, 113(4):700–765.

Bondy, A. G., Haefner, R. M., and Cumming, B. G. (2018). Feedback determines the structure of correlated variability in primary visual cortex. Nat. Neurosci., 21(4):598–606.

Britten, K. H., Newsome, W. T., Shadlen, M. N., Celebrini, S., and Movshon, J. A. (1996). A relationship between behavioral choice and the visual responses of neurons in macaque MT. Vis. Neurosci., 13:87–100.

Chicharro, D., Panzeri, S., and Haefner, R. M. (2017). Decision-related signals in the presence of nonzero signal stimuli, internal bias, and feedback. BioArxiv.

Choe, K. W., Blake, R., and Lee, S. H. (2014). Dissociation between neural signatures of stimulus and choice in population activity of human V1 during perceptual decision-making. J. Neurosci., 34(7):2725–2743.

Cicmil, N., Cumming, B. G., Parker, A. J., and Krug, K. (2015). Reward modulates the effect of visual cortical microstimulation on perceptual decisions. eLife, 4:e07832.

Cohen, M. R. and Maunsell, J. H. R. (2009). Attention improves performance primarily by reducing interneuronal correlations. Nat. Neurosci., 12(12):1594–1601.

Cohen, M. R. and Newsome, W. T. (2009). Estimates of the contribution of single neurons to perception depend on timescale and noise correlation. J. Neurosci., 29:6635–6648.

Cumming, B. G. and Nienborg, H. (2016). Feedforward and feedback sources of choice probability in neural population responses. Curr. Opinion Neurobiol., 37:126–132.

Ecker, A. S., Denfield, G. H., Bethge, M., and Tolias, A. S. (2016). On the structure of neuronal population activity under fluctuations in attentional state. J. Neurosci., 36(5):1775–1789.

Fetsch, C. R., Odean, N. N., Jeurissen, D., El-Shamayleh, Y., Horwitz, G. D., and Shadlen, M. N. (2018). Focal optogenetic suppression in macaque area MT biases direction discrimination and decision confidence, but only transiently. eLife, 7:e36523.

Fiser, J., Berkes, P., Orbán, G., and Lengyel, M. (2010). Statistically optimal perception and learning: from behavior to neural representations. Trends Cogn. Sci., 14:119–130.

Gold, J. I. and Shadlen, M. N. (2001). Neural computations that underlie decisions about sensory stimuli. Trends Cogn. Sci., 5:10–16.

Gold, J. I. and Shadlen, M. N. (2007). The neural basis of decision making. Annu. Rev. Neurosci., 30:535–574.

Goris, R. L., Movshon, J. A., and Simoncelli, E. P. (2014). Partitioning neuronal variability. Nat. Neurosci., 17:858–865.

Haefner, R. M. (2015). A note on choice and detect probabilities in the presence of choice bias. arXiv, page 1501.03173.

Haefner, R. M., Berkes, P., and Fiser, J. (2016). Perceptual decision-making as probabilistic inference by neural sampling. Neuron, 90(3):649–660.

Haefner, R. M., Gerwinn, S., Macke, J. H., and Bethge, M. (2013). Inferring decoding strategies from choice probabilities in the presence of correlated variability. Nat. Neurosci., 16:235–242.

Hanley, J. A. and McNeil, B. J. (1982). The meaning and use of the area under a receiver operating characteristic (ROC) curve. Radiology, 143(1):29–36.

Jasper, A. I., Tanabe, S., and Kohn, A. (2019). Predicting perceptual decisions using visual cortical population responses and choice history. J. Neurosci., 39(34):6714–6727.

Josić, K., Shea-Brown, E., Doiron, B., and de la Rocha, J. (2009). Stimulus-dependent correlations and population codes. Neural Comput., 21:2774–2804.

Kang, I. and Maunsell, J. H. (2012). Potential confounds in estimating trial-totrial correlations between neuronal response and behavior using choice probabilities. J. Neurophysiol., 108:3403–3415.

Kohn, A. and Smith, M. A. (2005). Stimulus dependence of neuronal correlation in primary visual cortex of the macaque. J. Neurosci., 25(14):3661–3673.

Krug, K., Curnow, T. L., and Parker, A. J. (2016). Defining the V5/MT neuronal pool for perceptual decisions in a visual stereo-motion task. Phil. Trans. R. Soc. B, 371:20150260.

Lakshminarasimhan, K. J., Pouget, A., DeAngelis, G. C., Angelaki, D. E., and Pitkow, X. (2018). Inferring decoding strategies for multiple correlated neural populations. PLoS Comput. Biol., 14(9):e1006371.

Lange, R. D. and Haefner, R. M. (2017). Characterizing and interpreting the influence of internal variables on sensory activity. Current Opinion in Neurobiology, 46:84–89.

Lee, D., Port, N. L., Kruse, W., and Georgopoulos, A. P. (1998). Variability and correlated noise in the discharge of neurons in motor and parietal areas of the primate cortex. J. Neurosci., 18:1161–1170.

Lee, T. S. and Mumford, D. (2003). Hierarchical bayesian inference in the visual cortex. J. Opt. Soc. Am. A Opt. Image Sci. Vis., 20:1434–1448.

Maunsell, J. H. and Treue, S. (2006). Feature-based attention in visual cortex. Trends Neurosci., 29:317–322.

Michelson, C., Pillow, J. W., and Seidemann, E. (2017). Majority of choice-related variability in perceptual decisions is present in early sensory cortex. BioArxiv.

Moreno-Bote, R., Beck, J., Kanitscheider, I., Pitkow, X., Latham, P., and Pouget, A. (2014). Information-limiting correlations. Nat. Neurosci., 17(10):1410–1417.

Nienborg, H., Cohen, M. R., and Cumming, B. G. (2012). Decision-related activity in sensory neurons: correlations among neurons and with behavior. Annu. Rev. Neurosci., 35:463–483.

Nienborg, H. and Cumming, B. G. (2006). Macaque V2 neurons, but not V1 neurons, show choice-related activity. J. Neurosci., 26:9567–9578.

Nienborg, H. and Cumming, B. G. (2009). Decision-related activity in sensory neurons reflects more than a neuron’s causal effect. Nature, 459:89–92.

Nienborg, H. and Cumming, B. G. (2010). Correlations between the activity of sensory neurons and behavior: how much do they tell us about a neuron’s causality? Current Opinion in Neurobiology, 20(3):1–6.

O’Connell, R. G., Shadlen, M. N., Wong-Lin, K., and Kelly, S. P. (2018). Bridging neural and computational viewpoints on perceptual decision-making. Trends Neurosci., 41(11):838–852.

Owen, D. B. (1956). Tables for computing bivariate normal probabilities. Annals of Mathematical Statistics, 27:1075–1090.

Park, I. M., Meister, M. L. R., Huk, A. C., and Pillow, J. W. (2014). Encoding and decoding in parietal cortex during sensorimotor decision-making. Nature Neurosci., 17(10):1395–1403.

Parker, A. J. and Newsome, W. T. (1998). Sense and the single neuron: probing the physiology of perception. Annu. Rev. Neurosci., 21:227–277.

Pillow, J. W. and Scott, J. G. (2012). Fully bayesian inference for neural models with negative-binomial spiking. Proceedings of the 25th Conference on Advances in Neural Information Processing Systems (NIPS 2012).

Pitkow, X., Liu, S., Angelaki, D. E., DeAngelis, G. C., and Pouget, A. (2015). How can single sensory neurons predict behavior? Neuron, 87(2):411–423.

Ponce-Alvarez, A., Thiele, A., Albright, T. D., Stoner, G. R., and Deco, G. (2013). Stimulus-dependent variability and noise correlations in cortical MT neurons. Proc Natl Acad Sci U S A, 110(32):13162–7.

Rao, R. P. and Ballard, D. H. (1999). Predictive coding in the visual cortex: a functional interpretation of some extra-classical receptive-field effects. Nat. Neurosci., 2(1):79–87.

Romo, R. and Salinas, E. (2003). Flutter discrimination: neural codes, perception, memory and decision making. Nat. Rev. Neurosci., 4(3):203–218.

Runyan, C. A., Piasini, E., Panzeri, S., and Harvey, C. D. (2017). Distinct timescales of population coding across cortex. Nature, 548(7665):92–96.

Sanayei, M., Chen, X., Chicharro, D., Distler, C., Panzeri, S., and Thiele, A. (2018). Perceptual learning of fine contrast discrimination changes neuronal tuning and population coding in macaque V4. Nat. Commun., 9(1):4238.

Shadlen, M. N., Britten, K. H., Newsome, W. T., and Movshon, J. A. (1996). A computational analysis of the relationship between neuronal and behavioral responses to visual motion. J. Neurosci., 16(4):1486–1510.

Shushruth, S., Mazurek, M., and Shadlen, M. N. (2018). Comparison of decision-related signals in sensory and motor preparatory responses of neurons in area LIP. J. Neurosci., 38(28):6350–6365.

Siegel, M., Buschman, T. J., and Miller, E. K. (2015). Cortical information flow during flexible sensorimotor decisions. Science, 348:1352–1355.

Smolyanskaya, A., Haefner, R. M., Lomber, S. G., and Born, R. T. (2015). A modality-specific feedforward component of choice-related activity in MT. Neuron, 87(1):208–219.

Steinmetz, N. A., Zatka-Haas, P., Carandini, M., and Harris, K. D. (2019). Distributed coding of choice, action and engagement across the mouse brain. Nature.

Tajima, C. I., Tajima, S., Koida, K., Komatsu, H., Aihara, K., and Suzuki, H. (2016). Population code dynamics in categorical perception. Scientific Reports, 6:22536.

Thielscher, A. and Pessoa, L. (2007). Neural correlates of perceptual choice and decision making during fear-disgust discrimination. J. Neurosci., 27(11):2908–2917.

Truccolo, W., Eden, U. T., Fellows, M. R., Donoghue, J. P., and Brown, E. N. (2005). A point process framework for relating neural spiking activity to spiking history, neural ensemble, and extrinsic covariate effects. J. Neurophysiol., 93:1074–1089.

Tsunada, J., Cohen, Y. E., and Gold, J. I. (2019). Post-decision processing in primate prefrontal cortex influences subsequent choices on an auditory decision-making task. eLife, 8:e46770.

Tsunada, J., Liu, A. S., Gold, J. I., and Cohen, Y. E. (2016). Causal contribution of primate auditory cortex to auditory perceptual decision-making. Nat. Neurosci., 19(1):135–142.

Urai, A. E., de Gee, J. W., Tsetsos, K., and Donner, T. H. (2019). Choice history biases subsequent evidence accumulation. eLife, 8:e46331.

van Vugt, B., Dagnino, B., Vartak, D., Safaai, H., Panzeri, S., Dehaene, S., and Roelfsema, P. R. (2018). The threshold for conscious report: Signal loss and response bias in visual and frontal cortex. Science, 360:537–542.

Wimmer, K., Compte, A., Roxin, A., Peixoto, D., Renart, A., and de la Rocha, J. (2015). Sensory integration dynamics in a hierarchical network explains choice probabilities in cortical area MT. Nature Communications, 6:6177.

Yang, H., Kwon, S. E., Severson, K. S., and OConnor, D. H. (2016). Origins of choice-related activity in mouse somatosensory cortex. Nat. Neurosci., 19(1):127–134.

Yu, X. and Gu, Y. (2018). Probing sensory readout via combined choice-correlation measures and microstimulation perturbation. Neuron, 100:715–727.

Zaidel, A., DeAngelis, G. C., and Angelaki, D. E. (2017). Decoupled choice-driven and stimulus-related activity in parietal neurons may be misrepresented by choice probabilities. Nat. Commun., 8(1):715.

## References

Arnold, B. C. and Beaver, R. J. (2000). Hidden truncation models. The Indian Journal of Statistics, 62:23–35.

Azzalini, A. (2005). The skew-normal distribution and related multivariate families. Scandinavian Journal of Statistics, 32(2):159–188.

Carnevale, F., de Lafuente, V., Romo, R., and Parga, N. (2013). An optimal decision population code that accounts for correlated variability unambiguously predicts a subject’s choice. Neuron, 80(6):1532–1543.

Dayan, P. and Abbot, L. F. (2001). Theoretical Neuroscience, Computational and Mathematical Modeling of Neural Systems. The MIT press, Cambridge, Massachusetts.

